# *Pde5a* Deficiency Prevents Diet-Induced Obesity via Adipose cAMP-PKA Activation Enhancing Fat Browning

**DOI:** 10.1101/2025.02.05.636635

**Authors:** F. Campolo, O. Giampaoli, F. Barbagallo, B. Palmisano, A. Di Maio, F. Sciarra, F. Rizzo, S. Monti, S. Cardarelli, M.R. Assenza, E. Poggiogalle, A. Patriarca, F. Sciubba, A. Filippini, A. Lenzi, D. Gianfrilli, M. Giorgi, S. Dolci, F. Naro, M. Sampaolesi, M. Riminucci, M. Mancini, A. Miccheli, L. Tessarollo, M.A Venneri, A.M Isidori

## Abstract

Cyclic nucleotides are critical regulators of adaptive thermogenesis and adipogenesis, with their intracellular levels finely tuned by phosphodiesterases. Phosphodiesterase type 5 (PDE5A) modulates cyclic guanosine monophosphate levels in adipocytes. While PDE5A inhibition has shown promise in patients with diabetes, its role in metabolism remains unclear. Using *Pde5a* knockout mouse models, we demonstrated that mice lacking *Pde5a* exhibit enhanced browning of white adipose tissue and reduced hepatic fat content. Following high-fat diet, *Pde5a*^−/−^ mice are resistant to obesity, displaying improved glucose metabolism and enhanced thermogenesis. These protective effects stem from an early developmental knockdown of *Pde5a*, leading to a metabolic reprogramming driven by cAMP-PKA pathway activation. The convergence of cGMP and cAMP signaling orchestrates thermogenic and systemic metabolic adaptations. Our findings establish PDE5A as a novel regulator of energy homeostasis, suggesting its inhibition as a valuable adjuvant therapy for metabolic disorders.

## INTRODUCTION

Adipose tissue (AT) has long been recognized as the principal site of energy storage, playing a critical role in whole-body energy homeostasis^1^. However, recent advances have transformed our understanding of AT, revealing it to be a dynamic endocrine organ composed of a heterogeneous population of cells including mature and immature adipocytes, immune cells, endothelial cells, progenitor and stem cells that collectively orchestrate complex metabolic responses. Dysregulation in the composition and function or excessive expansion of adipose tissue drives the complications observed in obesity and related conditions^2^. As the global prevalence of morbid obesity and metabolic-associated fatty liver disease (MAFLD) continue to rise^3^, the need to identify novel molecular targets and pathways governing energy metabolism becomes ever more urgent.

One promising but underexplored avenue is the pharmacological modulation of cyclic nucleotide signaling through the inhibition of phosphodiesterases (PDEs), enzymes responsible for the hydrolysis of cyclic nucleotides. Among these, phosphodiesterase 5 (PDE5A) stands out as the most extensively studied and clinically relevant, owing to the availability of human safe and affordable inhibitors. PDE5A selectively hydrolyzes cyclic guanosine monophosphate (cGMP)^4^, and its pharmacological inhibition has been shown to induce UCP1 expression, promote thermogenic activity, and stimulate the browning of subcutaneous white adipose tissue^5,6^. Additionally, PDE5A inhibition has been implicated in adipogenesis, as evidenced by increased intracellular lipid droplet accumulation and the upregulation of key adipogenic markers during 3T3-L1 preadipocyte differentiation^7^.

Preclinical and clinical studies seem to suggest that PDE5A inhibitors (PDE5i) improve insulin resistance and glucose metabolism highlighting their therapeutic potential for metabolic disorders^8,9^. We have extensively tested PDE5i in type 2 diabetes mellitus ^10–14^; however, the precise mechanisms underlying these effects remains poorly understood. Building on these findings, we aim to dissect the specific contributions of PDE5A in the regulation of glucose homeostasis and lipid metabolism. To achieve this purpose, we have developed global and conditional *Pde5a* knockout mouse models and we have investigated their metabolic phenotype, fat distribution and functional properties under basal conditions and in response to thermogenic and dietary challenges.

## RESULTS

The cGMP-Protein Kinase G (PKG) signaling pathway plays a crucial role in brown adipocyte differentiation, with several evidences demonstrating that pharmacological enhancement of cGMP in adipocytes induce UCP1 expression and drive the thermogenic program^5,6,15^. cGMP intracellular levels are primarily regulated by the *Pde5a* and *Pde9a* isoenzymes, both of which are expressed in mature adipocytes^9,16,17^. While recent evidence has shown that *Pde9a* is involved in the cold-induced thermogenic program of brown adipose tissue (BAT)^16^, the role of *Pde5a* in this process warrants further investigation. To determine whether PDE5A contributed to white adipose tissue (WAT) browning and thermogenesis, we generated a *Pde5a* knockout mouse model (*Pde5a^-/-^)* and analyzed its adipose tissue under basal conditions and after thermogenic stimuli.

### *Pde5a* ablation boosts browning of WAT and *in vivo* thermogenesis

*Pde5a^-/-^* mice are viable, fertile, have a normal life span, and were born at normal Mendelian ratios. Gross morphological inspection revealed a healthy phenotype, with no overt abnormalities and an apparently normal behavior, suggesting that loss of *Pde5a* does not affect mouse development and viability (**Figure S1C**). We measured cGMP-phosphodiesterase activity in the absence and presence of the selective PDE5A inhibitor sildenafil in mouse embryonic fibroblasts (MEF) cultures derived from *Pde5a*^-/-^ and wild-type mice. The analysis revealed that both the *Pde5a* ablation and pharmacological inhibition resulted in a comparable reduction of cGMP-phosphodiesterase activity (**Figure S1D**). Moreover, the addition of sildenafil to *Pde5a^-/-^* MEFs did not further reduce cGMP hydrolysis, confirming that sildenafil’s primary target in adipocytes is the PDE5A. To test whether

*Pde5*a ablation increases cGMP levels and activates the PKG signaling pathway, we analyzed the phosphorylation status of vasodilator stimulated phosphoprotein (VASP) in the same cell extracts. We found that global *Pde5a* deficiency significantly increased PKG-dependent phosphorylation of VASP on Serine^2^^39^ indicating heightened PKG activity (**Figure S1E**).

While histomorphometric analyses of adipose tissue depots showed no major differences in body weight or fat mass under basal condition between *Pde5a^-/-^*and wild-type mice (**Figure 1A-C)**, a significant reduction of adipocyte area was observed in both white and brown adipocytes of *Pde5a^-/-^* mice (**Figure 1D-E**). This reduction in adipocyte size is characteristic of ‘WAT-browning’, a process associated with increased non-shivering thermogenesis and metabolic activity^18^. Uncoupling protein 1 (UCP1) is a key player in the process by which BAT increases energy expenditure^19^. To investigate whether WAT browning occurs in *Pde5a^-/-^*mice, we analyzed the expression of canonical BAT markers. Notably, *Ucp1* expression was significantly upregulated in all adipose tissue depots of *Pde5a^-/-^* mice, suggesting that the absence of *Pde5a* promotes the “beigeing” of WAT while also fostering the thermogenic program in BAT (**Figure 2A-B**). In support of this observation, we found an overall increase in the expression of other thermogenic or BAT-enriched genes, including Peroxisome proliferator-activated receptor γ (*Ppar*γ), PR domain containing 16 (*Prdm16*), CCAAT/enhancer-binding protein a (*Cebp*α) and CCAAT/enhancer-binding protein b (*Cebp*β), Fibroblast growth factor 21 (*Fgf21*) and Pyruvate Dehydrogenase Kinase 4 (*Pdk4*) in *Pde5a^-/-^* WAT, suggesting that *Pde5a* ablation is accompanied by an increase in ‘*beige’* adipocytes within WAT (**Figure 2C**).

**Figure 1.**
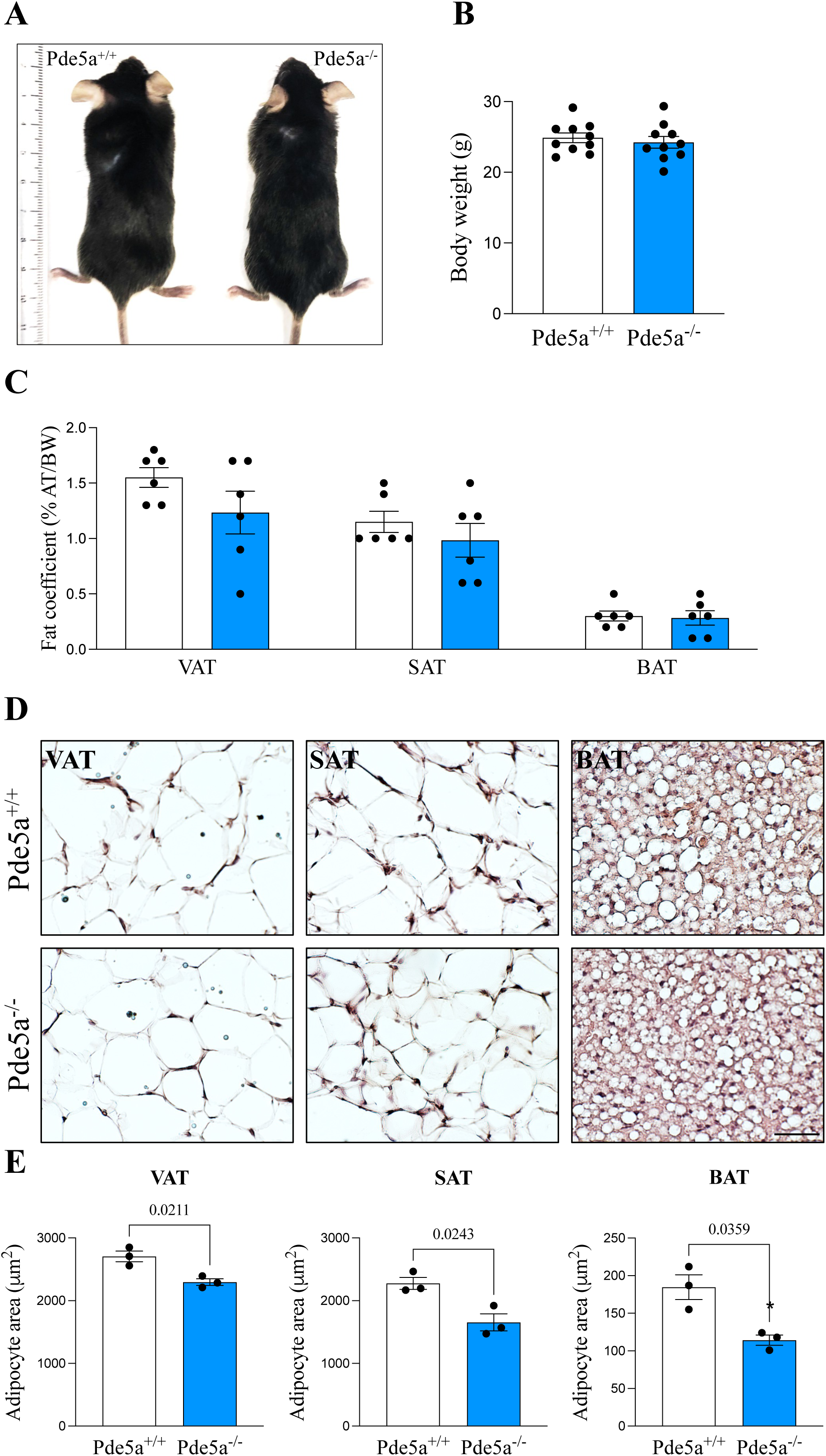
White adipose tissue from *Pde5a^-/-^* mice displays morphological features of brown fat. (A) Representative view of 2-month-old *Pde5a^+/+^* and *Pde5a^-/-^* male mice. (B) Body weight of *Pde5a^+/+^* (*n =10*) and *Pde5a^-/-^* (*n =10*) mice fed a normal diet. (C) Adipose tissue (epididymal visceral adipose tissue VAT, inguinal subcutaneous adipose tissue SAT, interscapular brown adipose tissue BAT) weight as a percentage of total body weight of 2-month-old *Pde5a^+/+^* (*n =6, white*) and *Pde5a^-/-^* (*n =6, blue*) male mice. (D) H&E staining of adipose tissue sections of VAT, SAT and BAT from 2-month-old *Pde5a^+/+^* and *Pde5a^-/-^* male mice; scale bar = 50 μm. (E) Average area of adipocytes (100 cells/mouse) in FFPE sections obtained from VAT, SAT and BAT of 2-month-old *Pde5a^+/+^* (*n =3, white*) and *Pde5a^-/-^* (*n =3, blue*) male mice. Results in scatter dot plot graphs are shown as mean ± SEM. Statistical analysis was performed using Student t-test.

**Figure 2.**
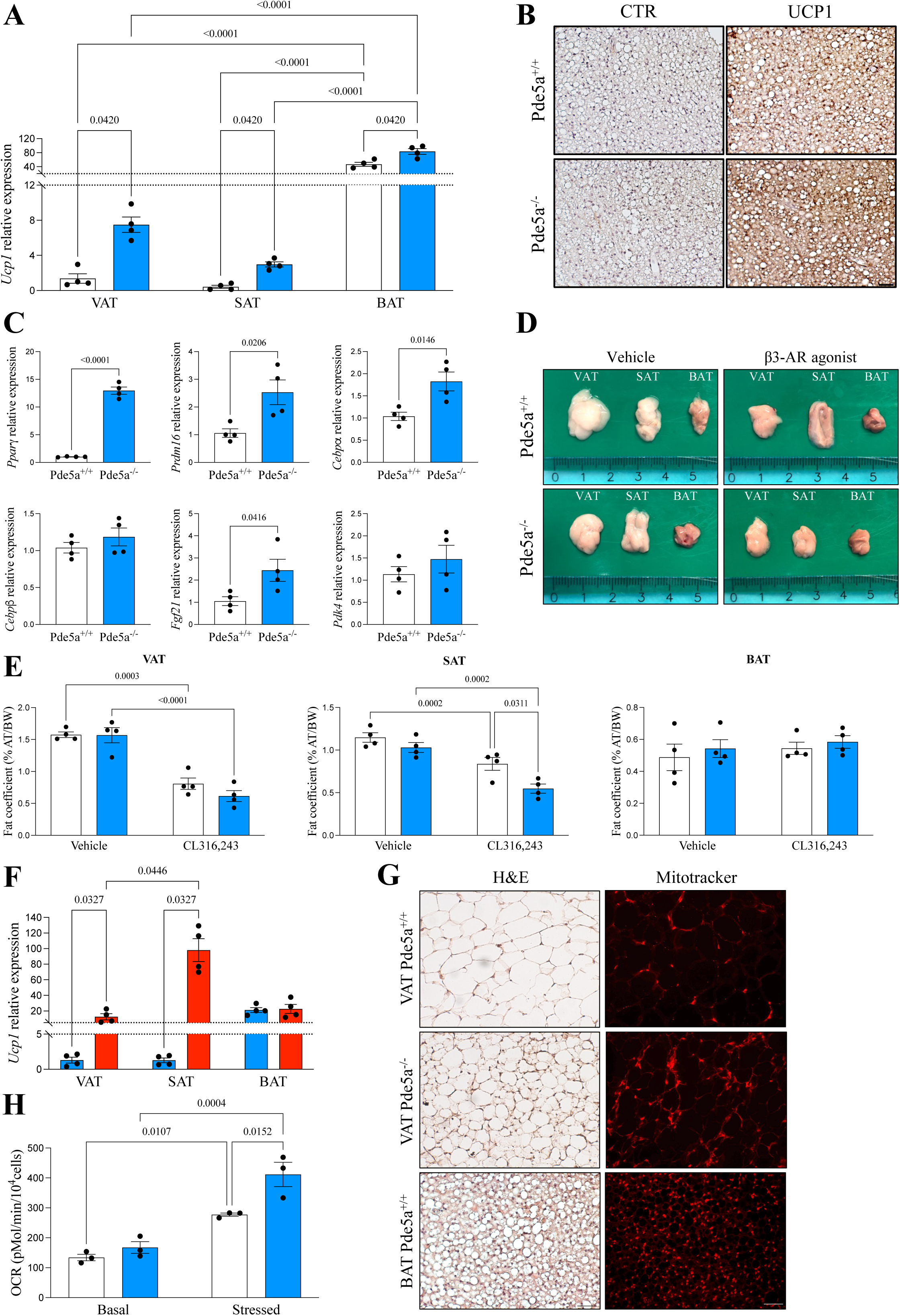
*Pde5a* ablation induces WAT browning and enhances thermogenesis. (A) qPCR gene expression analysis of *Ucp1* performed on adipose tissue depots obtained from *Pde5a^+/+^* (*n =4, white*) and *Pde5a^-/-^* (*n =4, blue*) mice. *Hprt1* was used as housekeeping gene for normalization. Data are presented as dot plots with column bars ± SEM showing all the experimental samples. Statistical analysis was performed using two-way ANOVA test. (B) Immunohistochemistry staining of BAT sections of *Pde5a^+/+^* (upper panels) and *Pde5a^-/-^* (lower panels) mice showing increased UCP1 expression in *Pde5a^-/-^* mice as compared to wild-type mice. Left panels = unstained sections; right panels = UCP1 stained sections. Scale bar = 100 μm. (C) qPCR gene expression analysis of thermogenic markers and BAT-enriched genes performed on VAT obtained from *Pde5a^+/+^* (*n =4, white*) and *Pde5a^-/-^* (*n =4, grey*) mice. *Hprt1* was used as housekeeping gene for normalization. Data are presented as dot plots with column bars ± SEM. Statistical analysis was performed using Student t-test. (D) Representative fat pads images of *Pde5a^+/+^* (upper panels) and *Pde5a^-/-^* (lower panels) mice following saline (left panels, vehicle) or CL316,243 (right panels, β3-AR agonist) injection. (E) Adipose tissue depots weight as a percentage of total body weight of *Pde5a^+/+^* (*n =4, white*) and *Pde5a^-/-^* (*n =4, blue*) mice following saline or CL316,243 treatment. Data are presented as dot plots with column bars ± SEM. Statistical analysis was performed using two-way ANOVA test. (F) qPCR gene expression analysis of *Ucp1* performed on adipose tissue depots obtained from *Pde5a^-/-^* mice following saline (*n =4, blue*) and CL316,243 (*n =4, red*) treatment. *Hprt1* was used as housekeeping gene for normalization. Data are presented as dot plots with column bars ± SEM. Statistical analysis was performed using two-way ANOVA test. (G) Hematoxylin and Eosin (H&E) staining and Mitotracker staining from wild-type and *Pde5a*^-/-^ VAT and wild-type BAT. Scale bar = 50 μm. (H) Respiration and glycolysis analysis on VAT cultures from *Pde5a^+/+^* (*n =3, white*) and *Pde5a^-/-^* (*n =3, blue*) mice trough quantification of basal and stressed Oxygen consumption rate (OCR). Data are presented as dot plots with column bars ± SEM. Statistical analysis was performed using Student t-test.

To further assess the impact of *Pde5a* deletion on fat thermogenic capacity *in vivo*, *Pde5a^-/-^* mice were treated with a β3-adrenergic receptor (β3-AR) selective agonist (CL316,243) for ten days. Macroscopic analysis of tissues from vehicle and CL316,243-treated mice revealed that fat depots from *Pde5a^-/-^* mice appeared constitutively browner in color compared to wild-type controls, resembling a thermogenic activated phenotype (**Figure 2D**). Following β3-AR agonist treatment, we found a significant reduction in visceral fat (VAT) in both *Pde5a^-/-^* and wild-type mice. However, this effect was not observed in the subcutaneous fat (SAT), the depot most prone to browning. In SAT, the absence of *Pde5a* resulted in a greater reduction in fat mass following adrenergic stimulation compared to wild-type mice (**Figure 2E**). This effect was accompanied by a robust increase in *Ucp1* expression in SAT from CL316,243 treated *Pde5a^-/-^*mice (**Figure 2F**), suggesting a pivotal role of PDE5A in regulating SAT plasticity.

Mitochondrial dynamics play a critical role in energy metabolism and glucose homeostasis^20^. To further elucidate the mechanism underlying the increased thermogenic potential in *Pde5a* knockout mice, we examined mitochondria density and function in AT. Mitotracker staining of VAT revealed increased mitochondrial density in *Pde5a^-/-^* mice compared to wild-type controls, likely attributable to the presence of a fair amount of multilocular adipocytes within *Pde5a* deficient WAT (**Figure 2G**). To verify whether the increased number of mitochondria also reflects a greater mitochondrial activity, we analyzed mitochondrial function in primary adipocytes. Oxygen consumption rate (OCR), a measure of mitochondrial aerobic respiration, and extracellular acidification rate (ECAR)^21^, an indicator of lactate production, were both increased in primary adipocytes derived from *Pde5a^-/-^*mice compared to wild-type controls (**Figure 2H and S2A**). This increase reached statistical significance following the addition of test compounds (Oligomycin and FCCP), which allowed for the estimation of maximal respiratory capacity. Collectively, these data suggest that *Pde5a* deficiency does not broadly alter mitochondrial function across all adipocytes but rather promotes the accumulation of *beige* adipocytes within *Pde5a* deficient WAT. The increase in *beige* adipocytes drives the enhanced thermogenic activity, indicating that the primary role of PDE5A relies in regulating adipose tissue plasticity, rather than directly mitochondrial biogenesis. This observation is further supported by the analysis of Peroxisome proliferator-activated receptor-gamma coactivator 1 alpha (PGC1α), whose mRNA expression levels are not significantly different in VAT from *Pde5a* ko mice compared to wild-type (**Figure S2B)**.

The increased thermogenic capacity prompted us to investigate surface body temperature using infrared thermography that accurately reflects changes in BAT activity *in vivo*^22^. We measured surface body temperature of mice under thermoneutral conditions (23°C) and following cold-induced thermogenesis (4°C). Infrared thermocamera images and surface skin temperature analysis showed that, under thermoneutral conditions, *Pde5a* ko mice display a significantly higher interscapular skin temperature, as well as greater difference between interscapular and lumbar back skin temperature compared to wild-type mice (**Figure 3A**). The same trend can be observed during cold exposure, although statistical significance was reached only after normalization for the lumbar surface temperature, as expected. This is consistent with the observation that interscapular BAT in the wild-type appropriately responds to cold challenge, while BAT in the *Pde5a* deficient mice is already basally more activated, even under thermoneutral conditions (**Figure 3B**).

**Figure 3.**
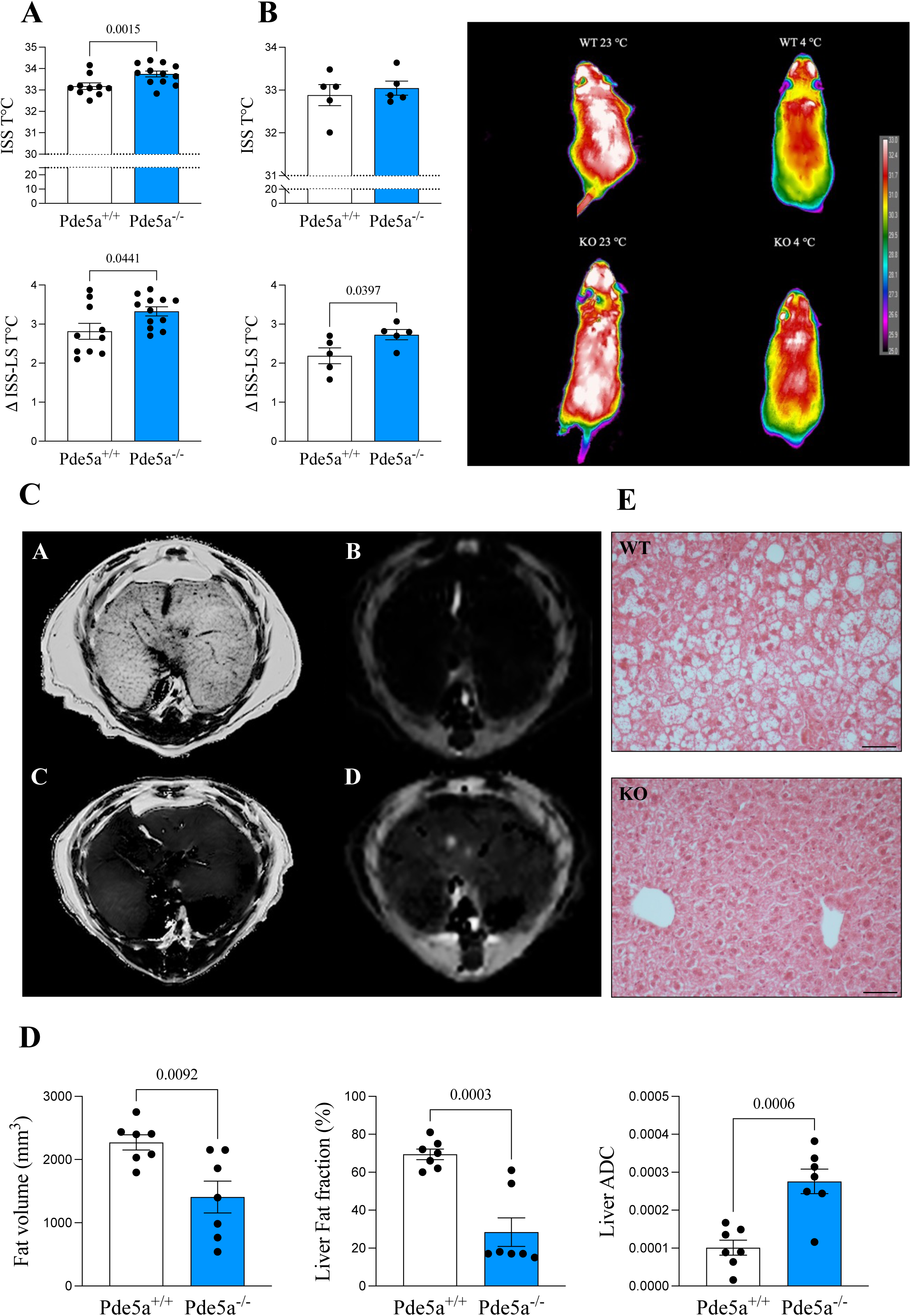
*Pde5a* deficiency increases energy expenditure and thermogenesis in vivo. (A-B) Quantification of average interscapular surface skin temperature (ISS, *top*) and delta interscapular skin surface temperature and lumbar surface temperature (ISS-LS, *bottom*) on *n= 10 Pde5a^+/+^* mice and *n = 12 Pde5a*^-/-^ mice/each group at 23°C (A) and 4°C (B). Representative infrared imaging of *Pde5a*^+/+^ (*right upper panels*) and *Pde5a*^-/-^ (*right lower panels*) mice under thermoneutral condition (23°C) or after cold exposure (4°C). Data are presented as dot plots with column bars ± SEM. Statistical analysis was performed applying one-way ANOVA test for upper panels and ANCOVA test comparing post-exposure temperature values and accounting for baseline temperature values as covariate (lower panels). (C) Exemplificative liver Fat Fraction (*A, C panels*) and Apparent Diffusion Coefficient (*B, D panels*) MRI maps of *Pde5a*^+/+^ (*A, B panels*) and *Pde5a*^-/-^ (*C, D panels*). (D) Dot plot analysis of subcutaneous and visceral fat volume (*left graph*), liver fat fraction (*middle graph*) and liver apparent diffusion coefficient (*right graph*). Results in scatter dot plot graphs are shown as mean ± SEM (n=7 for each group). Statistical analysis was performed using Student t-test. (E) H&E staining of liver sections from 12 months old *Pde5a*^+/+^ and *Pde5a*^-/-^ male mice; scale bar = 50μm.

In summary, these data demonstrate that constitutive *Pde5a* ablation induces WAT browning and enhances thermogenesis *in vivo*, suggesting that PDE5A plays a pivotal role in modulating the thermogenic program.

### *Pde5a* deficiency prevents aging-related liver steatosis and diet-induced obesity

To determine the long-term effect of having an enhanced browning *in vivo*, we performed abdominal Magnetic Resonance Imaging (MRI) in 12-month-old mice. Imaging analysis clearly showed that *Pde5a^-/-^* mice display reduced visceral and subcutaneous fat volumes compared to controls (**Figure 3C**). Notably, a striking difference in liver Fat Fraction (FF) and Apparent Diffusion Coefficient (ADC) was observed, with *Pde5a* deficient mice showing markedly lower levels (**Figure 3D**). Subsequent histological analysis of liver sections confirmed protection against aging-induced liver steatosis, as demonstrated by the absence of hepatocyte ballooning and inflammatory infiltrates in the liver parenchyma (**Figure 3E**).

Having established that *Pde5a* deficiency may confer protection against age-related metabolic alterations, we next investigated whether also protects against diet-induced metabolic challenges.

First, *Pde5a* deficient mice were subjected to either a low-fat normal chow (NC) or high-fat diet (HFD) for 12 weeks, following the experimental protocol outlined in **Figure S2C**. Mice on a normal diet showed no significant changes in body weight. However, when on an HFD, body mass increased dramatically in wild-type but much less in *Pde5a* knockout mice, starting from the fourth week of dietary change (**Figure 4A-B**).

**Figure 4.**
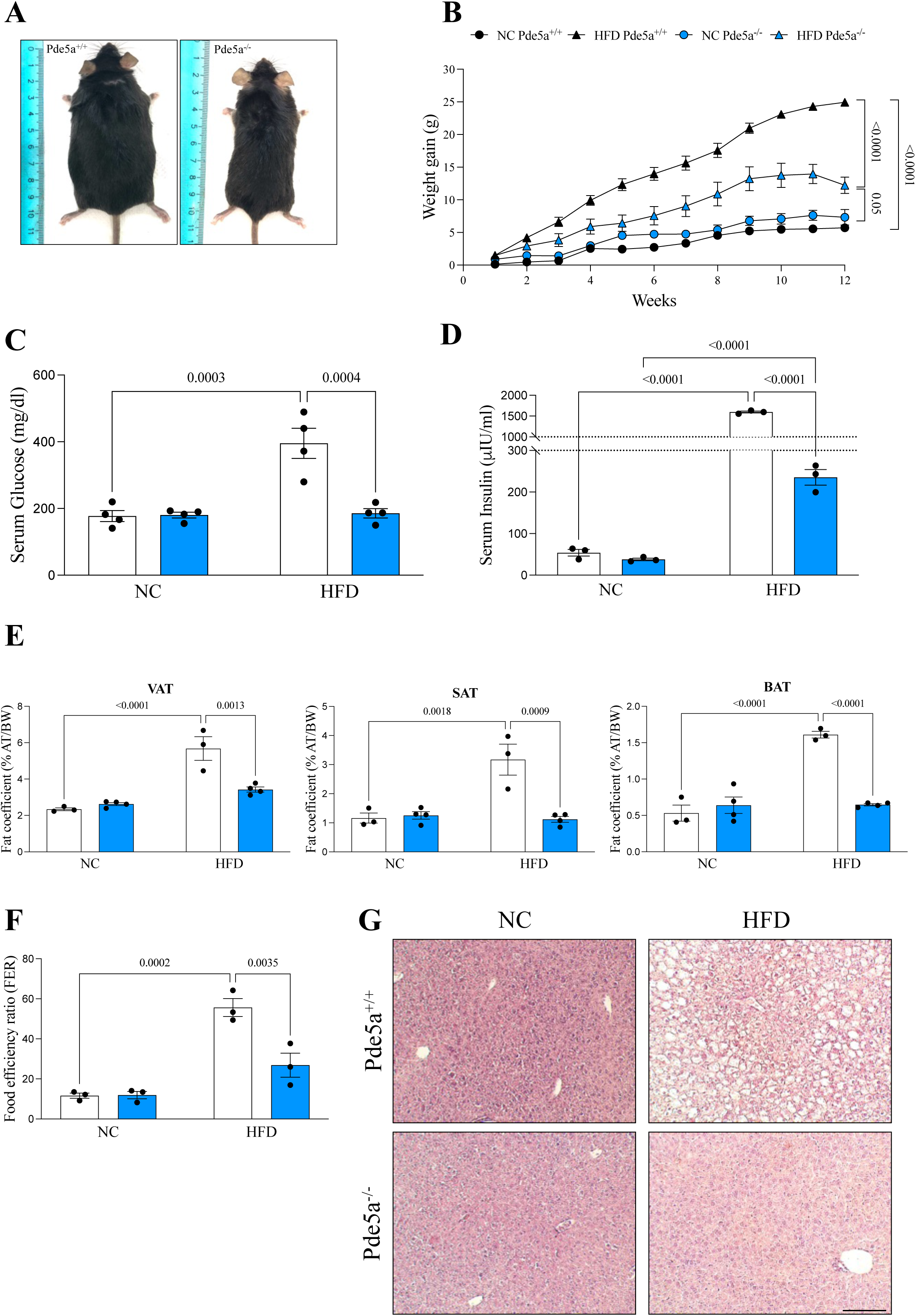
*Pde5a* deletion elevates energy expenditure and confers protection from HFD-induced obesity. (A) Dorsal view of *Pde5a*^+/+^ and *Pde5a*^-/-^ male mice after 12 weeks feeding with high fat diet (HFD) at age of 6-month-old. (B) Relative change in body weight (weight gain) of *Pde5a*^-/-^ and their control littermates fed with HFD or normal chow (NC) over 12 weeks, showing onset of obesity in wild-type but not in Pde5^-/-^ mice (n= 9 each group). Data are expressed as mean ± SEM. Statistical analysis was performed applying a three-way ANOVA test. (C-D) Measurements of fasting serum glucose (C) and insulin (D) levels in *Pde5a*^-/-^ (*blue*) and WT (*white*) mice after 12 weeks feeding with NC and HFD (n = 4). Data are presented as dot plots with column bars ± SEM. Statistical analysis was performed using two-way ANOVA test. (E) Adipose tissue depots weight as a percentage of total body weight of *Pde5a*^+/+^ (*white*) and *Pde5a*^-/-^ (*blue*) mice after 12-weeks feeding with NC and HFD (n=4 each group). Data are presented as dot plots with column bars ± SEM. Statistical analysis was performed using two-way ANOVA test. (F) Food efficiency ratio (FER) percentage of weight gain/food intake in *Pde5a*^-/-^ (*blue*) and wt (*white*) mice after 12 weeks feeding with NC (n=10) and HFD (n = 10). Data are presented as dot plots with column bars ± SEM. Statistical analysis was performed using two-way ANOVA test. (G) H&E staining of liver from *Pde5a*^+/+^ (*top*) and *Pde5a*^-/-^ (*bottom*) mice after 12-weeks feeding with NC (*left panels*) and HFD (*right panels*). Scale bar =100μm.

Second, HFD is typically associated with, dyslipidemia, peripheral insulin resistance and glucose intolerance^23^. Remarkably, *Pde5a* deficient mice were resistant to the hyperglycemia and hyperinsulinemia induced by the HFD (**Figure 4C-D)**. Consistent with these findings, HFD-fed *Pde5a^-/-^* mice displayed a significant reduction of fat mass compared to HFD-fed wild-type mice, which aligned with the differences observed in main adipose tissue depots (**Figure 4E**). Importantly, these differences were not attributable to variations in food intake, as daily food consumption was comparable between *Pde5a^-/-^*and wild-type mice under both NC or HFD conditions (**Figure S2D**). Additionally, no significant difference in physical activity, measured by total distance traveled, were observed between the two groups (**Figure S2E**). Conversely, the food efficiency ratio (FER), calculated as the ratio of weight gain to food intake, which was similar in both genotypes under NC, was significantly lower in *Pde5a* deficient mice under HFD, indicating improved metabolic efficiency (**Figure 4F**).

Third, HFD feeding is known to cause liver damage due to lipid accumulation and systemic chronic low-grade inflammation^24,25^. In agreement with the observed metabolic findings, HFD-fed *Pde5a^-/-^* mice were protected from HFD-induced liver steatosis (**Figure 4G**) and exhibited lower serum levels of pro-inflammatory cytokines IL1α and INFy (**Figure S2F-G**).

Together, these findings demonstrate that constitutive deletion of *Pde5a* protects mice from both aging and diet-induced obesity, while mitigating age- and HFD-induced hepatic steatosis and inflammation.

### *Pde5a* ablation affects glucose homeostasis, insulin and adipokines

To further investigate the metabolic phenotype associated to *Pde5a* deficiency, we assessed whole-body glucose homeostasis by measuring plasma glucose levels and insulin sensitivity. Both the Glucose Tolerance Test (GTT) and Insulin Tolerance Test (ITT) showed that, while fasting glucose levels were similar between the two groups (**Figure 4C**), the early glucose response (15-30 min) following glucose administration was unexpectedly higher in *Pde5a^-/-^* than wild-type mice, before returning to similar levels at 60 min and thereafter. Interestingly, insulin levels were nearly identical at all time points. Since the initial response to glucose infusion is typically associated with a stop of hepatic glucose outflow, the fact that the fasting, 60 and 120 min glucose levels, as well as insulin at all time points, were identical between groups, suggest that *Pde5a^-/-^* mice exhibit a normal insulin sensitivity, but altered hepatic glucose output (**Figure 5A-B**). Thus, the transient difference in early response to glucose infusion seems mainly due to a more efficient reduction in liver glucose output in the wild-type, as confirmed by metabolomic analysis.

**Figure 5.**
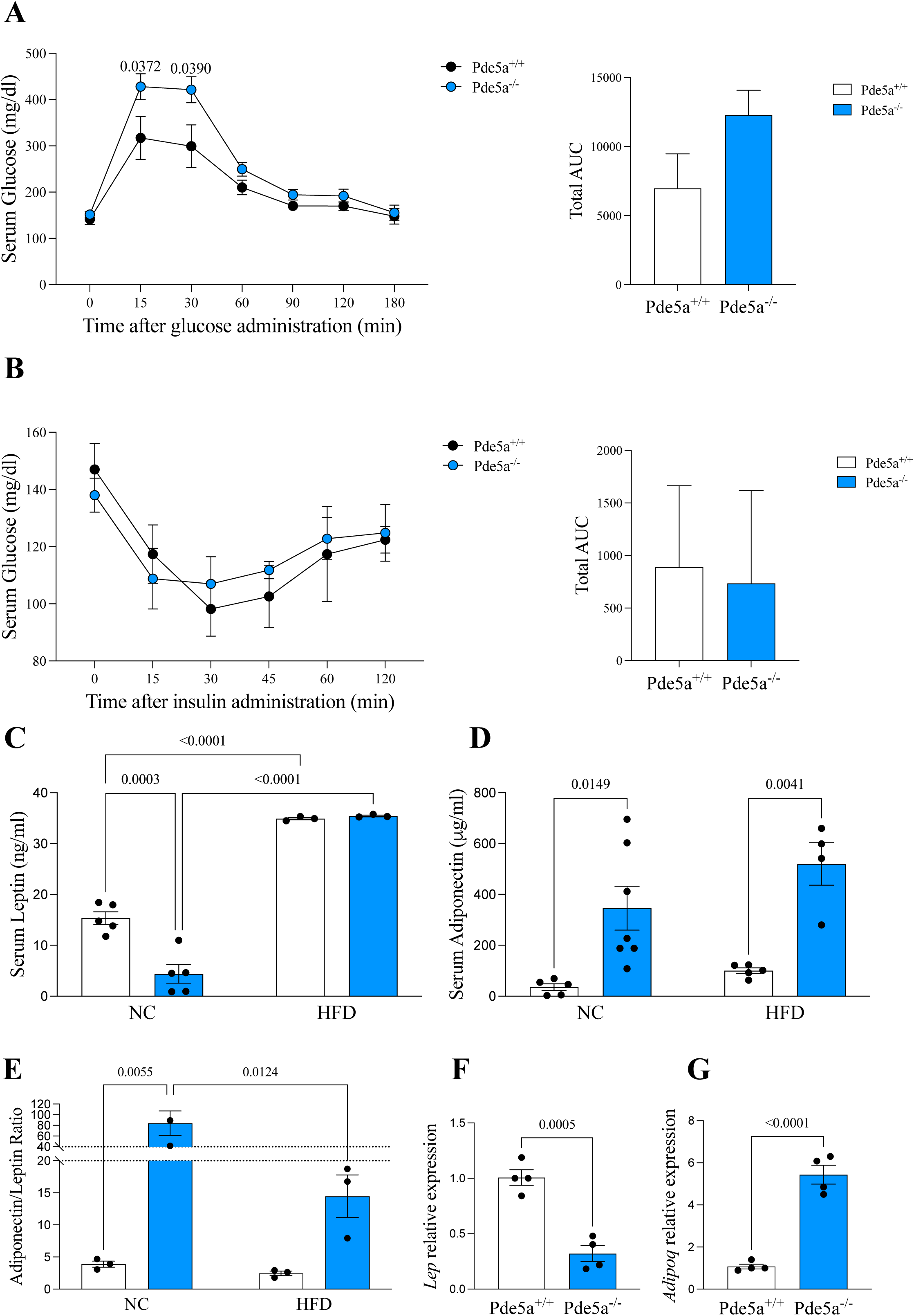
*Pde5a* deficiency confers a constitutive metabolic healthy phenotype to mice. (A-B) Intraperitoneal glucose tolerance test (IP-GTT, A) and intraperitoneal insulin tolerance test (IP-ITT, B) in 2-month-old *Pde5a*^+/+^ (*white*) and *Pde5a*^−/−^ (*blue*) male mice (n=5/group). Area under the curve (AUC) of data in A and B are shown on right panels. Statistical analysis was performed using Student t-test. (C) Measurements of serum leptin levels in *Pde5a*^-/-^ (*blue*) and wild-type (*white*) mice after 12 weeks feeding with NC (n=5) and HFD (n = 3). Data are presented as dot plots with column bars ± SEM. Statistical analysis was performed using two-way ANOVA test. (D) Measurements of serum adiponectin levels in *Pde5a*^-/-^ (*blue*) and wild-type (*white*) mice after 12 weeks feeding with NC (n=5) and HFD (n = 4). Data are presented as dot plots with column bars ± SEM. Statistical analysis was performed using two-way ANOVA test. (E) Measurement of Adiponectin/Leptin ratio in *Pde5a*^-/-^ (*blue*) and wild-type (*white*) mice after 12 weeks feeding with NC (n=5) and HFD (n = 4). Data are presented as dot plots with column bars ± SEM. Statistical analysis was performed using two-way ANOVA test. (F-G) qPCR gene expression analysis of *Lep* (F) and *AdipoQ* (G) performed on visceral adipose tissue obtained from *Pde5a*^+/+^ (*n =4, white*) and *Pde5a*^-/-^ (*n =4, blue*) mice. *Hprt1* was used as housekeeping gene for normalization. Data are presented as dot plots with column bars ± SEM. Statistical analysis was performed using Student t-test.

To complete the metabolic characterization of these mice, we also measured adipokines, which interact dynamically with insulin to promote metabolic homeostasis^26^. Under physiological conditions, serum levels of adiponectin and leptin are inversely correlated, with obesity typically associated with elevated leptin and decreased adiponectin levels^27^. *Pde5a^-/-^* mice displayed significantly lower levels of leptin (**Figure 5C**) and higher levels of adiponectin (**Figure 5D**), resulting in a marked reduction in the adiponectin/leptin ratio (**Figure 5E**), a surrogate marker of metabolic impairment. These findings are supported by the corresponding changes in the expression of *Lep* and *Adipoq* genes in WAT (**Figure 5F-G**).

The mitigated weight gain in *Pde5a* knockout mice under HFD was accompanied by significantly lower serum insulin and glucose levels, alongside higher levels of adiponectin. Surprisingly, however, there was no difference in plasma leptin levels, suggesting a dissociation between adipocyte size area, fat depots mass, and adipokine expression. Once again, the data suggests a shift in adipose tissue plasticity triggered by *Pde5a* deficiency.

### *Pde5a* knockout triggers organ-specific reprogramming of glucose usage and fatty acids metabolism

We employed nuclear magnetic resonance (NMR) based metabolomic to analyze glucose-responsive tissues from starved mice and after glucose administration (metabolic challenge). Fasting data did not reveal striking differences between the two genotypes (data not shown), consistently with the grossly normal phenotype of *Pde5a^-/-^* under basal conditions. Distinct metabolic responses were, however, evident under metabolic challenge. Resonance assignment for each matrix for both for ^1^H and ^13^C nuclei are provided in **Table 1**. The use of [1,2,3,4,5,6-^13^C_6_] glucose in conjunction with 1^13^C-NMR spectroscopy enabled the labeling of intermediates to investigate the metabolic pathways involved in glucose utilization. ^13^C labeling was applied to liver, kidney, VAT, BAT and serum. However, labeled isotopomers were detected exclusively in the liver and kidney, likely due to the lower concentration of labeled metabolites in other tissues, which hindered a robust resonance detection (data not shown).

**Table 1.**
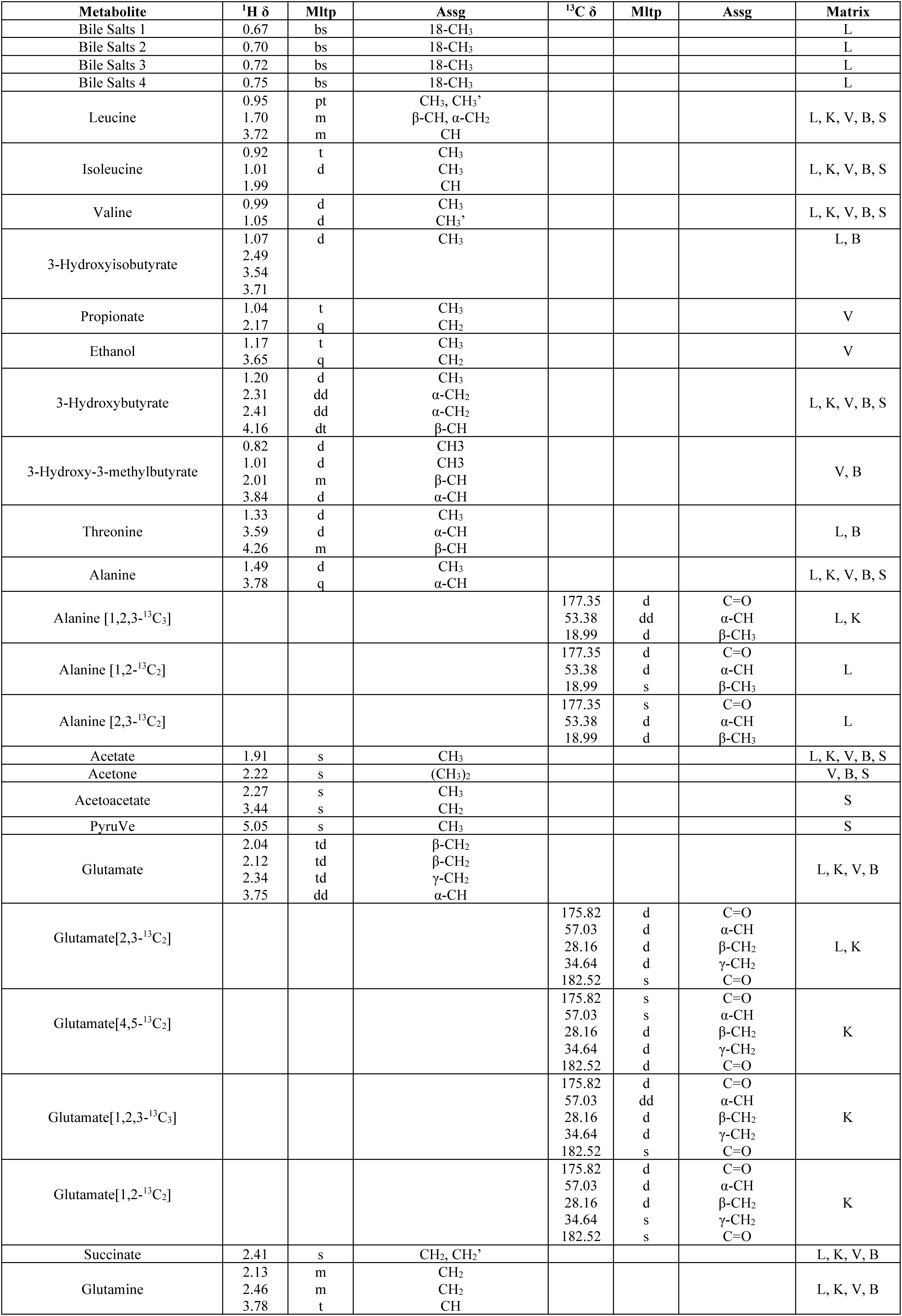
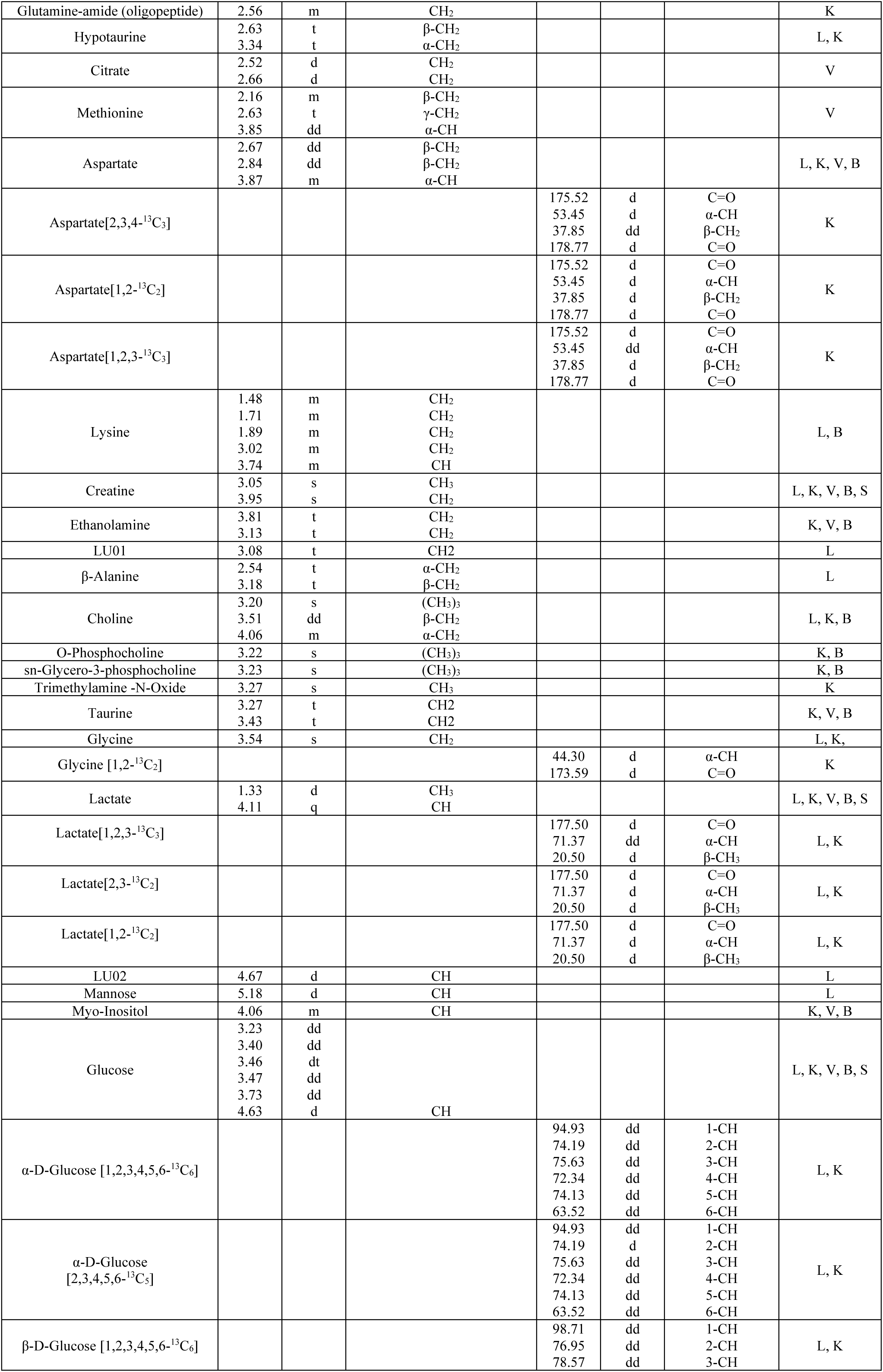
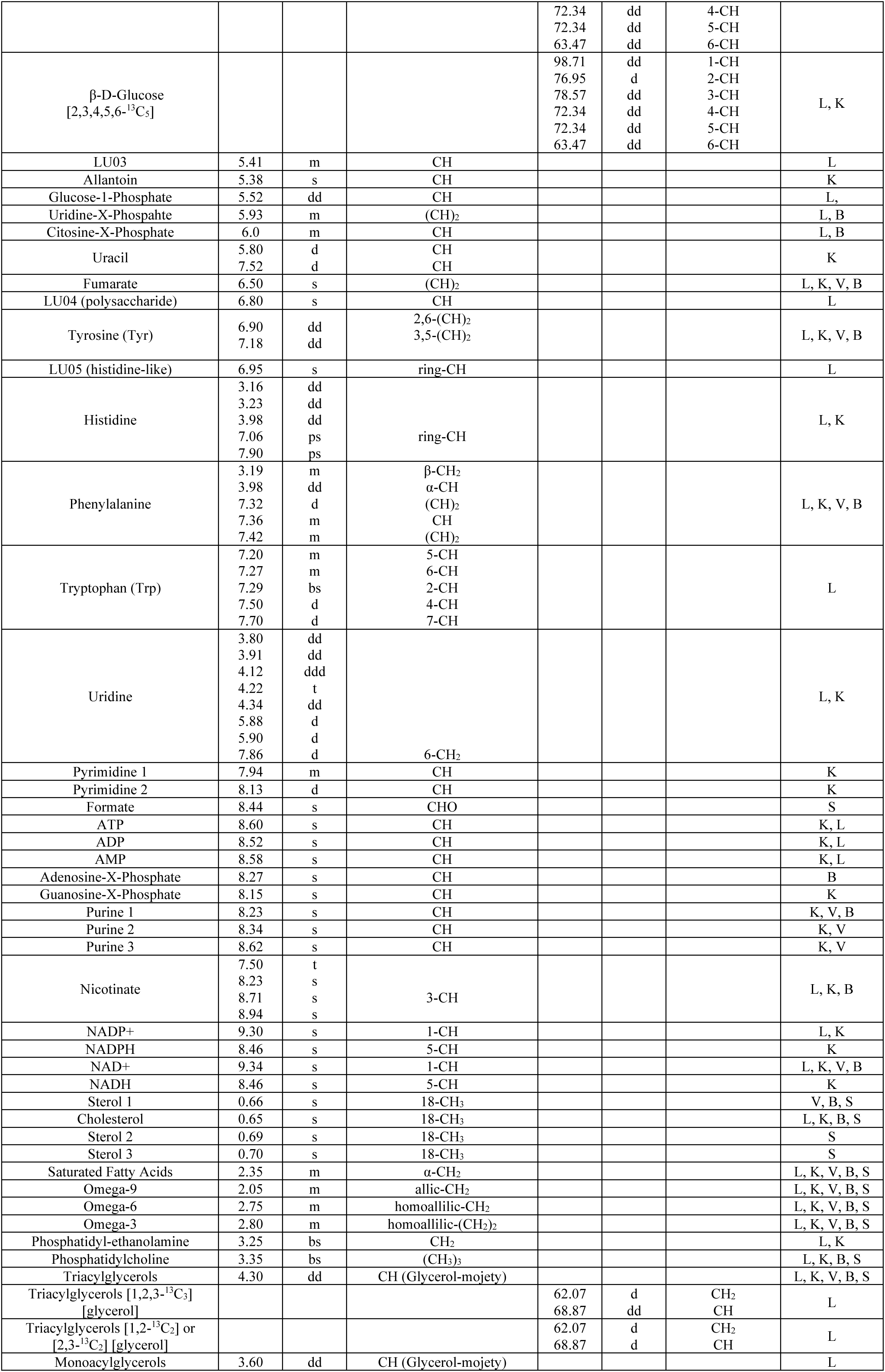
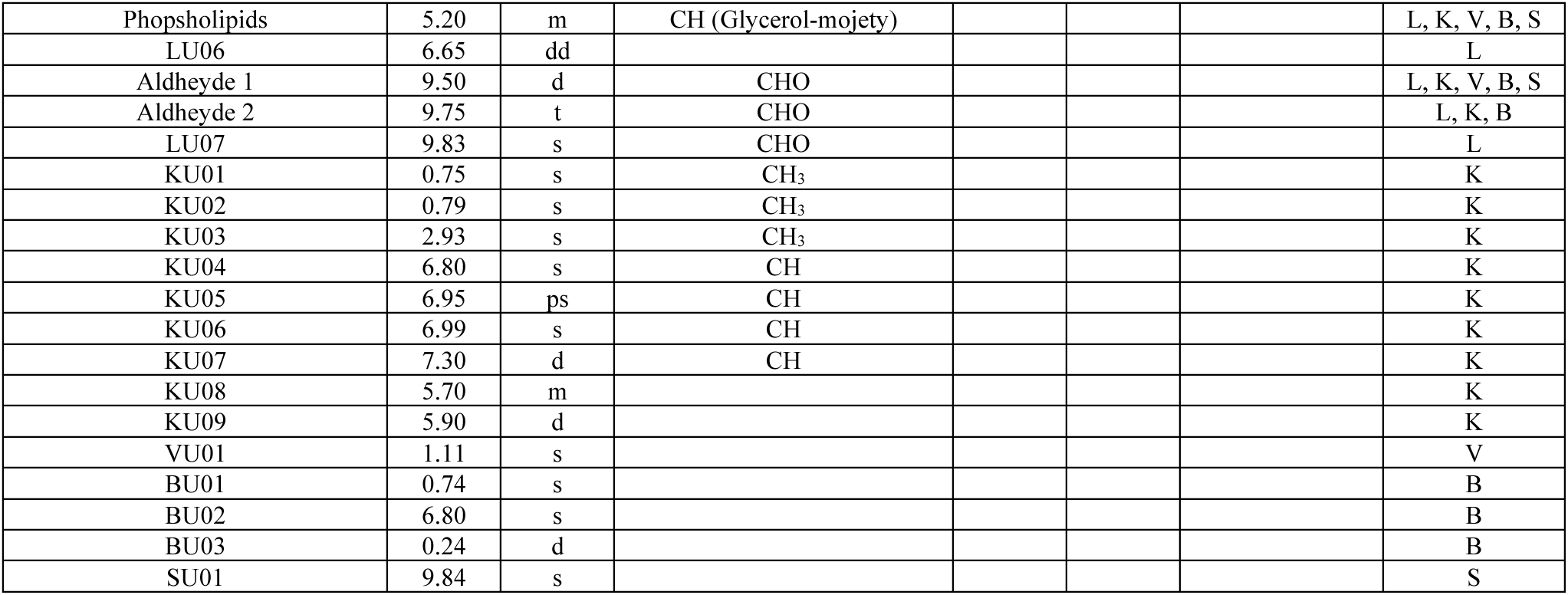
^1^H and ^13^C resonance assignment for the metabolites observed through NMR spectroscopy. *s: singlet; bs: broad singlet; d: doublet; dd: double doublet; t: triplet; q: quadruplet; td: triplet pf doublets; m: multiplet; mltp: multiplicity; assg: assignment; U: unknown compound; K: kidney; L: liver; V: VAT; B: BAT; S: serum*.

The ^1^H-NMR liver spectra after glucose administration produced a PLS-DA model with strong prediction accuracy (91%) and precision (94%). Notably, bile salts and glucose-1-phosphate levels were significantly higher in *Pde5a^-/-^* mice, while levels of long-chain fatty acids, triacylglycerols, organic acids, lactate, choline, amino acids, and glucose were lower. Regression coefficient analysis also revealed that Pde*5a* deficiency leads to lower levels of 3-hydroxybutyrate, a ketone body generated by β-hydroxybutyrate dehydrogenase activity in the ketogenetic pathway (**Figure 6A**).

**Figure 6.**
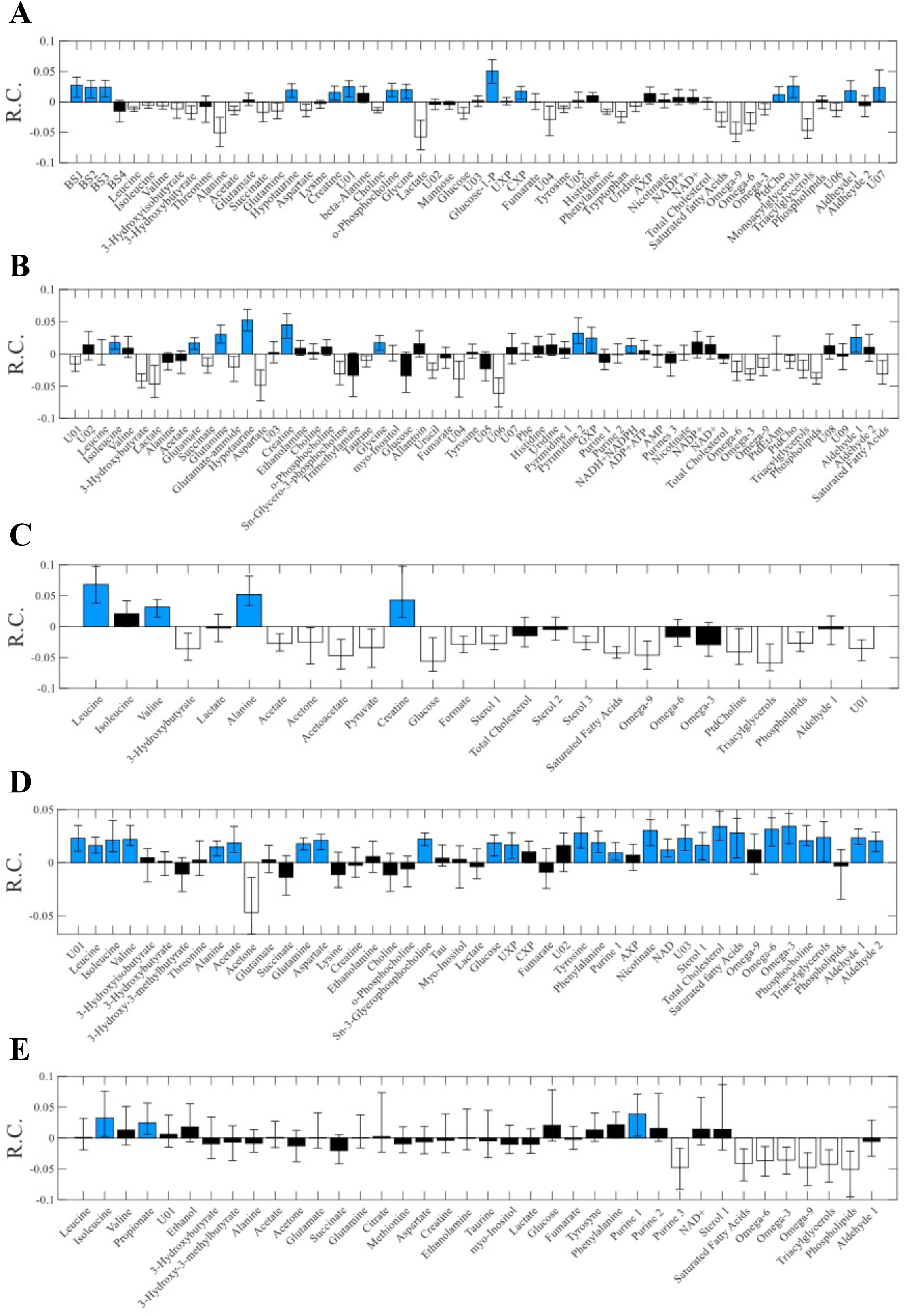
NMR reveals metabolic pathways constitutively altered in *Pde5a* deficient mice. Panel for the PLS-DA model of insulin sensitive tissues and serum obtained from Pde5 knockout and wild-type mice after glucose administration. Regression coefficient values are shown for liver (A), kidney (B), serum (C), BAT (D) and VAT (E). Variables significantly higher in *Pde5a* ko are depicted in *blue*, variables significantly lower in *Pde5a* deficient mice are depicted in *white* and non-significantly modulated variables are shown in *black*.

The ^13^C-NMR liver spectra displayed complex patterns, documenting the presence of various isotopologue species (**Table 1**). From ^13^C-NMR liver spectra, 11 different labeled metabolites were identified. Metabolites derived from glycolysis revealed that *Pde5a^-/-^*mice exhibited significantly lower levels of lactate and glycerol moiety of triglyceride isotopomers compared to wild-type mice (**Table 2 and Figure S3A**). Notably, we did not observe the presence of labeled glutamate [4,5-^13^C_2_] in either *Pde5a^+/+^*and *Pde5a^-/-^* mice, suggesting that the Acetyl-CoA involved in the tricarboxylic acid (TCA) cycle is primarily derived from β-oxidation of fatty acids rather than glycolysis. In addition, the glucose isotopomers ratios indicate that gluconeogenesis did not vary between genotypes, as evidenced by similar production of [2,3,4,5,6-^13^C_5_] glucose. Overall, liver metabolomics data showed a reduction in lactate and alanine concentrations, supporting a shift in glucose flux towards glycogen synthesis, rather than glycolysis. This fits with the higher glucose-1-phosphate observed in ^1^H-NMR experiment, and lower ^13^C labelling in glycolysis intermediates, as well as a reduced glycerol moiety labeling in triglycerides. The preferential storage of glucose into glycogen of *Pde5a-/-* mice, also explains their lower glucose liver output that accounts for the differential early serum response during glucose infusion test.

**Table 2.** Fractional enrichments of quantified ^13^C labelled metabolites’ isotopomers for liver and kidney samples assessed by ^13^C NMR spectroscopy after Glucose [1,2,3,4,5,6-^13^C_6_] oral administration. Data are expressed as mean ± standard deviation. The ^13^C labelled glucose ratio of glucose [1,2,3,4,5,6-^13^C_6_] and [2,3,4,5,6-^13^C_5_] for liver and kidney samples obtained from *Pde5a^+/+^* and *Pde5a^-/-^* mice is also reported.

| Matrix | Metabolite | *Pde5a*<sup>+/+</sup> | *Pde5a*<sup>-/-</sup> | p value |
| --- | --- | --- | --- | --- |
| Liver | Alanine [1,2,3-<sup>13</sup>C<sub>3</sub>] | 10.17 ± 3.16 | 6.42 ± 2.21 | 0.142 |
|  | Alanine [1,2-<sup>13</sup>C<sub>2</sub>] | 1.41 ± 0.58 | 1.002 ± 0.52 | 0.403 |
|  | Alanine [2,3-<sup>13</sup>C<sub>2</sub>] | 2.76 ± 0.65 | 1.98 ± 0.30 | 0.111 |
|  | Glutamate [2,3-<sup>13</sup>C<sub>2</sub>] | 3.16 ± 1.07 | 1.91 ± 0.59 | 0.126 |
|  | Lactate [1,2,3-<sup>13</sup>C<sub>3</sub>] | 12.29 ± 1.38 | 7.15 ± 2.45 | **0.019** |
|  | Lactate [2,3-<sup>13</sup>C<sub>2</sub>] | 2.29 ± 0.16 | 1.48 ± 0.23 | **0.003** |
|  | Lactate [1,2-<sup>13</sup>C<sub>2</sub>] | 0.73 ± 0.11 | 0.51 ± 0.05 | **0.025** |
|  | Triacylglycerols [1,2,3-<sup>13</sup>C<sub>3</sub>] [glycerol] | 1.08 ± 0.15 | 0.54 ± 0.24 | **0.016** |
|  | Triacylglycerols [1,2-<sup>13</sup>C<sub>2</sub>] or [2,3-<sup>13</sup>C<sub>2</sub>] [glycerol] | 0.77 ± 0.18 | 0.63 ± 0.15 | **0.033** |
| Kidney | Alanine [1,2,3-<sup>13</sup>C<sub>3</sub>] | 6,157 ± 2,148 | 9,676 ± 7,487 | 0.464 |
|  | Aspartate [2,3,4-<sup>13</sup>C<sub>3</sub>] | 1,384 ± 0,202 | 0,984 ± 0,099 | **0.021** |
|  | Aspartate [1,2-<sup>13</sup>C<sub>2</sub>] | 0,87 ± 0,222 | 0,471 ± 0,27 | 0.095 |
|  | Aspartate [1,2,3-<sup>13</sup>C<sub>3</sub>] | 0,778 ± 0,292 | 2,634 ± 1,005 | **0.022** |
|  | Glycine [1,2-<sup>13</sup>C<sub>2</sub>] | 0,733 ± 0,103 | 0,409 ± 0,277 | 0.107 |
|  | Glutamate [4,5-<sup>13</sup>C<sub>2</sub>] | 0,713 ± 0,154 | 0,438 ± 0,109 | **0.045** |
|  | Glutamate [1,2,3-<sup>13</sup>C<sub>3</sub>] | 1,147 ± 0,482 | 1,382 ± 1,257 | 0.772 |
|  | Glutamate [1,2-<sup>13</sup>C<sub>2</sub>] | 0,716 ± 0,115 | 0,656 ± 0,193 | 0.659 |
|  | Glutamate [2,3-<sup>13</sup>C<sub>2</sub>] | 1,895 ± 0,214 | 1,238 ± 0,415 | 0.051 |
|  | Lactate [1,2,3-<sup>13</sup>C<sub>3</sub>] | 11,277 ± 2,635 | 7,491 ± 1,384 | 0.070 |
|  | Lactate [2,3-<sup>13</sup>C<sub>2</sub>] | 2,016 ± 0,262 | 1,357 ± 0,196 | **0.013** |
|  | Lactate [1,2-<sup>13</sup>C<sub>2</sub>] | 0,638 ± 0,236 | 0,628 ± 0,416 | 0.973 |
| Matrix | <sup>13</sup>C labelled glucose ratio | *Pde5a*<sup>+/+</sup> | *Pde5a*<sup>-/-</sup> | p value |
| Liver | C2/C1 | 1.20 ± 0.03 | 1.25 ± 0.07 | 0.247 |
| Kidney | C2/C1 | 1.70 ± 0.20 | 1.34 ± 0.11 | **0.041** |

In the kidney, ^1^H-NMR spectra highlighted 62 metabolites (**Table 1**). The PLS-DA model showed 82% accuracy and 78% precision, with significant higher levels of glutamine, hypotaurine, creatine, pyrimidine compound, and GTP in *Pde5a^-/-^* mice (**Figure 6B**). Conversely, aspartate, 3-hydroxybutyrate, lactate, lipids and phospholipids were lower in *Pde5a^-/-^* mice after metabolic challenge. ^13^C-NMR data from kidney samples revealed 14 labeled intermediates. The presence of labeled glutamate [4,5-^13^C_2_] indicates the incorporation of labeled Acetyl-CoA derived from the glycolytic pathway, in contrast to what is observed in the liver. Yet, the fact that of glutamate [4,5-^13^C_2_] production is still reduced in the kidney of *Pde5a^-/-^* mice, confirms the observation made in the liver of a globally reduced glucose [1,2,3,4,5,6-^13^C_6_] flux through the glycolytic pathway (**Figure S3B and Table 2**), and the use of amino acids as alternative energy source.

Adipose tissue NMR analysis, complemented the findings observed in the liver and kidney. The PLS-DA model for BAT following glucose administration demonstrated a clear separation between wild-type and *Pde5a^-/-^* mice (92% accuracy and 88% precision). Most significant metabolites, including amino acids, organic acids, bile salts (BS1), glycerophosphocholine, phosphocholine, glucose, NAD, and lipids were positively correlated with the *Pde5a^-/-^* genotype, except for acetone, which was negatively correlated (**Figure 6D**). Similarly, the PLS-DA model for VAT indicated reduced levels of long-chain fatty acids and phospholipids in *Pde5a^-/-^* mice (**Figure 6E**).

Finally, serum NMR data after glucose administration recapitulates the metabolic shift observed in the other tissues, with a clear genotype separation (accuracy of 85% and precision of 85%) and a significant negative variation for lipids and ketone bodies, as inspected by PLS-DA model with a (**Figure 6C**).

Overall, the combined labeled (^13^C and ^1^H) and unlabeled NMR analyses suggest that Pde5a deficiency triggers significant metabolic reprogramming, affecting various tissues. This is particularly evident following metabolic challenges, especially in the liver, kidney, and BAT. These changes hint at an adaptive shift in glucose and lipid handling derived from *Pde5a*’s regulation of tissue-specific metabolism.

### *Pde5a* knockout mice display increased cAMP-PKA signaling in adipose tissue

The peculiar metabolic activation observed in BAT prompted us to explore the underlying molecular mechanism. Among these, the cAMP-PKA signaling pathway is considered crucial for browning phenotype and thermogenic activation ^21,28^. In adipocytes, cAMP levels are regulated by cGMP-inhibited phosphodiesterase 3 (PDE3), with the *Pde3b* isoform being highly expressed in adipose tissues^29^. Previous studies have shown that the ablation of *Pde3b* confers resistance against HFD, by promoting WAT browning and the activation of 5′-adenosine monophosphate (AMP)-activated protein kinase (AMPK) signaling^30,31,32^. Given this, we hypothesized that an interaction between cGMP and cAMP pathways might occur in the *beige* adipose tissue of *Pde5a^-/-^* mice. In our model, *Pde3b* mRNA levels were found significantly reduced (**Figure 7A**), leading to elevated cAMP levels (**Figure 7B**). This increase in cAMP was able to activate PKA, as evidenced by enhanced phosphorylation of PKA-substrates (**Figure 7C**), without compensatory change in the expression of other PDEs (**Figure S4A**). To test whether PDE5A stabilizes PDE3B protein, we treated *Pde5a*-overexpressing 3T3-L1 cells with cycloheximide (CHX). As shown in **Figure 7G**, PDE5A overexpression not only increases PDE3B basal expression but also delays its degradation. To further explore the PDE5A-PDE3B crosstalk, we performed *in vitro* silencing experiments in 3T3-L1 adipocytes. Silencing of *Pde3b* decreased *Pde5a* mRNA levels (**Figure 7D-E**), suggesting that *Pde3b* positively regulates *Pde5a* transcription. The downregulation of *Pde3b* observed in the *beige* adipose tissue AT from *Pde5a* ko mice appears to be mediated by increased levels of the transcriptional repressor ICER (inducible cAMP early repressor), which is activated in response to cAMP rise (**Figure 7F)**. Collectively these findings demonstrate that metabolic activation of *beige* AT observed in *Pde5a* ko mice results from the convergence between cAMP-PKA and cGMP-PKG signaling pathways.

**Figure 7.**
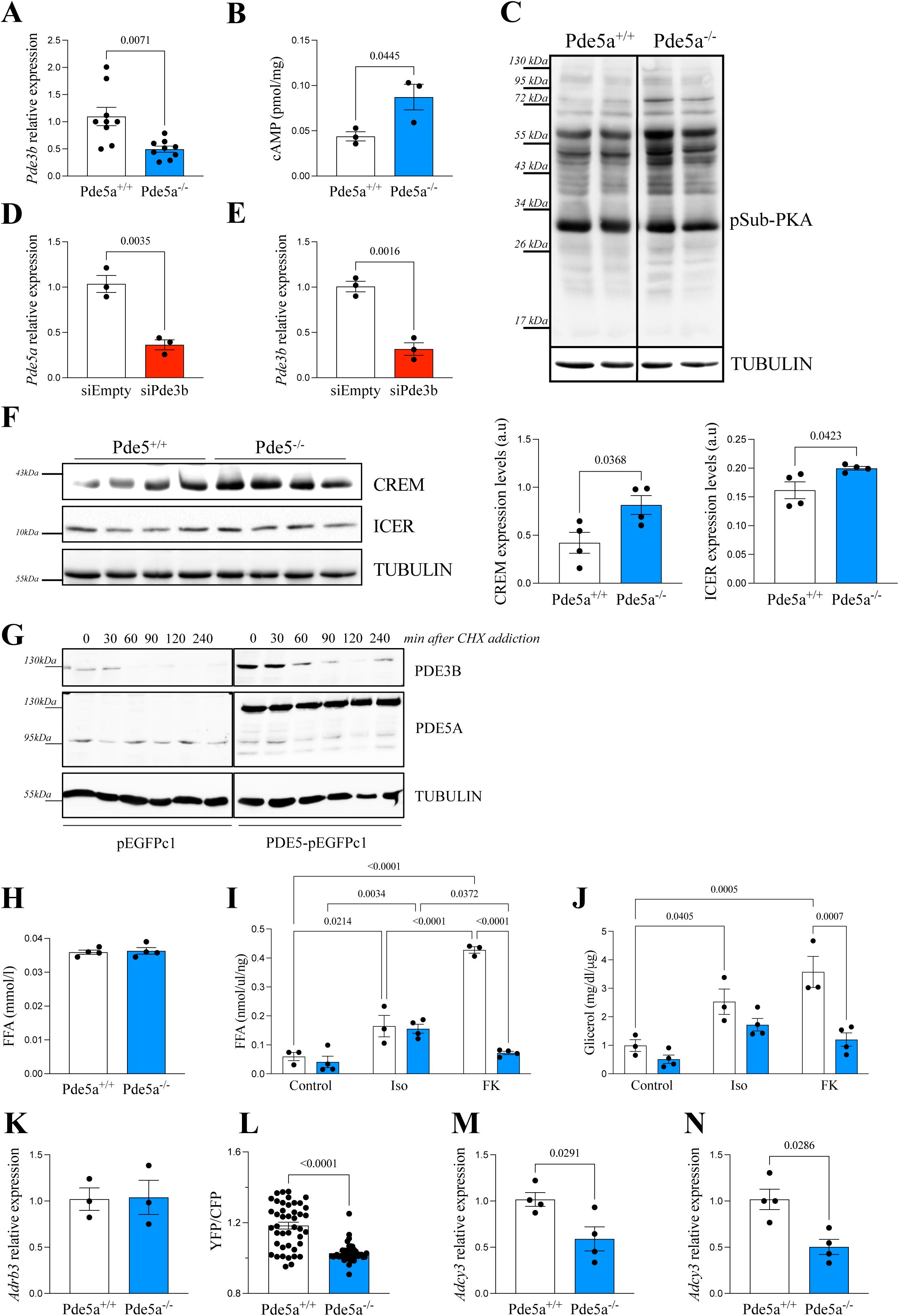
*Pde5a* knockout mice display increased cAMP-PKA signaling and impaired Adenylate Cyclase activity. (A) qPCR gene expression analysis of *Pde3b* performed on visceral adipose tissue obtained from *Pde5a*^+/+^ (*n =9, white*) and *Pde5a*^-/-^ (*n =9, blue*) mice. *Hprt1* was used as housekeeping gene for normalization. Data are presented as dot plots with column bars ± SEM. Statistical analysis was performed using Student t-test. (B) Quantification of intracellular cAMP levels through colorimetric assay in visceral adipose tissue lysates obtained from *Pde5a*^+/+^ (*n =3, white*) and *Pde5a*^-/-^ (*n =3, blue*) mice. Data are presented as dot plots with column bars ± SEM. Statistical analysis was performed using Student t-test. (C) Representative immunoblots of phosphorylated PKA substrate protein levels in whole lysates obtained from visceral adipose tissue obtained from *Pde5a*^+/+^ and *Pde5a*^-/-^ (*n =3*) mice. TUBULIN was used as housekeeping gene for normalization. (D-E) qPCR gene expression analysis of *Pde5a* (D) and *Pde3b* (E) performed on 3T3-L1 cells transfected with *siPde3b* or scramble control (siEmpty). (F) Immunoblots of CREM and ICER protein levels in whole lysates obtained from visceral adipose tissue obtained from *Pde5a*^+/+^ and *Pde5a*^-/-^ (*n =4*) mice. TUBULIN was used as housekeeping gene for normalization. Densitometric analyses for both targets are shown in the graphs on the right. Data are presented as dot plots with column bars ± SEM. Statistical analysis was performed using Student t-test. (G) PDE3B and PDE5A expression in 3T3-L1 cells transfected with GFP-PDE5A plasmid or empty GFP plasmid, treated with or without 5 mg/mL cycloheximide (CHX) for the indicated time (0-4 hours). Tubulin was used as housekeeping gene for normalization. (H-J) Basal and stimulated lipolysis evaluation trough serum free fatty acids quantification in *Pde5a^+/+^* and *Pde5a^-/-^* (*n=3*) mice (H). Free fatty acids (FFA, I) and glycerol (J) quantification in visceral adipocytes isolated from *Pde5a*^+/+^ and *Pde5a*^-/-^ (*n =4*) mice after lipolysis stimulation with 10 μM forskolin/isoproterenol. Data are presented as dot plots with column bars ± SEM. Statistical analysis was performed using Student t-test (H) or two-way ANOVA test (I, J). (K) qPCR gene expression analysis of *Adrb3* performed on visceral adipose tissue obtained from *Pde5a*^+/+^ (*n=3, white*) and *Pde5a*^-/-^ (*n=3, blue*) mice. *Hprt1* was used as housekeeping gene for normalization. Data are presented as dot plots with column bars ± SEM. (L) Normalized YFP/CFP FRET ratio on MEFs isolated from *Pde5a*^+/+^ and *Pde5a*^-/-^ mice transfected with AKAR3 PKA sensor and stimulated with 1μM forskolin. Data are presented as dot plots (n=50) with column bars ± SEM derived from three independent experiments. Statistical analysis was performed using Student t-test. (M-N) qPCR gene expression analysis of *Adcy3* performed on visceral adipose tissue (M) and MEFs (N) obtained from *Pde5a*^+/+^ (*n=4, white*) and *Pde5a*^-/-^ (*n=4, blue*) mice. *Hprt1* was used as housekeeping gene for normalization. Data are presented as dot plots with column bars ± SEM. Statistical analysis was performed using Student t-test.

### *Pde5a* deficiency impairs Adenylate Cyclase activity

Despite the augmented cAMP-PKA signaling, we observed lower free fatty acid (FFA) levels, glycerol moiety, and β-oxidation rate in *Pde5a* ko mice. This led us to examine lipolysis, a key process regulated by cAMP induced phosphorylation of perilipin and hormone sensitive lipase (HSL), which catalyzes triglycerides and diglycerides breakdown^8,29^, resulting in the release of FFA and glycerol^33,34^. Contrary to expectations, global deletion of *Pde5a* did not stimulate basal lipolysis (**Figure 7H-J**). Although *Pde5a* knockout VAT responded similarly to the β-ARs receptor activation with isoproterenol similar to wild-type, (**Figure 7K**), it failed to respond to forskolin, suggesting hypoactivation of adenylate cyclase (AC), the enzyme responsible for synthesizing cAMP from ATP. Using a fluorescence resonance energy transfer (FRET)-based sensor targeting PKA activity (AKAR3)^35^ we demonstrated that when overexpressed in MEFs from *Pde5a* ko mice and stimulated with isoproterenol, the PKA sensor exhibited similar FRET changes (data not shown). In contrast, MEF stimulation with forskolin resulted in a reduced FRET ratio, confirming the hypo-responsiveness of AC to forskolin observed in *Pde5a* deficient VAT (**Figure 7L**).

This observation prompted us to investigate AC expression in *Pde5a* ko mice. Notably, the AC3 isoform (encoded by *Adcy3*), the most abundantly expressed in VAT and MEFs, was significantly reduced (**Figure 7M-N**). In the *Pde5a^-/-^* mice, this compensatory mechanism may serve to counteract the constitutive increase of cAMP levels caused by *Pde3b* downregulation.

### Adipocyte-Specific *Pde5a* deletion is not sufficient to protect mice from HFD-induced obesity

The data so far indicate a metabolic reprogramming driven by adipose tissue *Pde5a*/*Pde3b* and AC-cAMP-PKA crosstalk. To determine at which stage this new equilibrium is established, we crossed mice harboring *LoxP*-flanked *Pde5a* alleles (*Pde5a^LoxP/LoxP^*) with transgenic mice expressing Adiponectin (Adipoq)-promoter driven Cre recombinase (*Adipoq^Cre^*) obtaining a mouse with committed adipocytes lacking *Pde5a* (*Pde5a*^Adpn-cKO^) (**Figure 8A and S4B**). The *Adipoq-Cre* recombination is restricted to mature adipocytes^36,37^, thus at a later stage of differentiation. These mice display only a 25% reduction of *Pde5a* mRNA levels extracted from VAT (**Figure 8B**) confirming the predominant expression of *Pde5a* in adipose endothelial cells (the VAT vascular fraction) compared to mature adipocytes^4,38,39^. A parallel reduction in *Pde3b* expression was detected in the same samples (**Figure 8C**), however, there were no differences in body weight when *Pde5a^Adpn-cKO^* mice were fed with HFD, suggesting that the mature adipocyte-specific loss of *Pde5a* is not sufficient to trigger metabolic reprogramming in these animals (**Figure 8D**).

**Figure 8.**
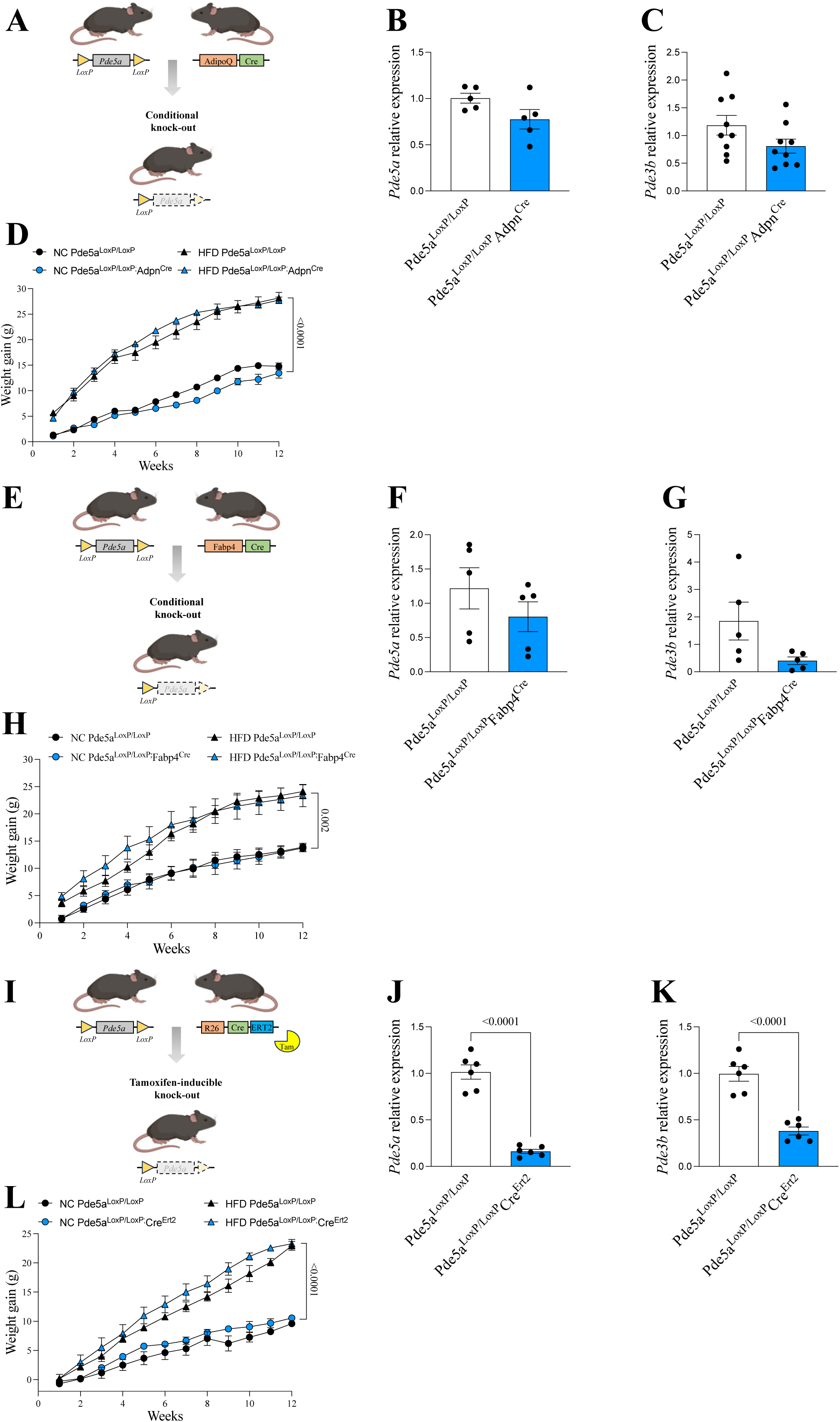
Adipocyte-specific *Pde5a* deletion did not protect mice from HFD-induced obesity. (A) Schematic illustration of targeting strategy for conditional knockout of *Pde5a* in mature adipocytes using Adpn^Cre^ mice. (B-C) qPCR gene expression analysis of *Pde5a* (B) and *Pde3b* (C) performed on visceral adipose tissue isolated from *Pde5a^LoxP/LoxP^* (*white, n=5*) and *Pde5a ^LoxP/LoxP^*; Adpn^Cre^ (*blue, n=5*). *Hprt1* was used as housekeeping gene for normalization. Data are presented as dot plots with column bars ± SEM. Statistical analysis was performed using Student t-test. (D) Relative change in body weight (weight gain) of *Pde5a^LoxP/LoxP^* (*white*) and *Pde5a ^LoxP/LoxP^; Adpn^Cre^* (*blue*) fed with HFD or normal chow (NC) over 12 weeks (n= 4 each group). Data are expressed as mean ± SEM. Statistical analysis was performed applying a three-way ANOVA test. (E) Schematic illustration of targeting strategy for conditional knockout of *Pde5a* in developing adipocytes using *Fabp4^Cr^*^e^ mice. (F-G) qPCR gene expression analysis of *Pde5a* (F) and *Pde3b* (G) performed on visceral adipose tissue isolated from *Pde5a^LoxP/LoxP^* (*white,* n=5) and *Pde5a ^LoxP/LoxP^*; *Fabp4^Cre^* (*blue,* n=5). *Hprt1* was used as housekeeping gene for normalization. Data are presented as dot plots with column bars ± SEM. Statistical analysis was performed using Student t-test. (H) Relative change in body weight (weight gain) of *Pde5a^LoxP/LoxP^* (*white*) and *Pde5a ^LoxP/LoxP^*; *Fabp4^Cre^ (blue*) fed with HFD or normal chow (NC) over 12 weeks (n= 6 each group). Data are expressed as mean ± SEM. Statistical analysis was performed applying a three-way ANOVA test. (I) Schematic illustration of targeting strategy for inducible ubiquitarian conditional knockout of *Pde5a* using *Cre^Ert2^* mice. (J-K) qPCR gene expression analysis of *Pde5a* (J) and *Pde3b* (K) performed on visceral adipose tissue isolated from *Pde5a^LoxP/LoxP^* (*white, n=5*) and *Pde5a ^LoxP/LoxP^*; *Cre^Ert2^* (*blue, n=5*). *Hprt1* was used as housekeeping gene for normalization. Data are presented as dot plots with column bars ± SEM. Statistical analysis was performed using Student t-test. (L) Relative change in body weight (weight gain) of *Pde5a^LoxP/LoxP^* (*white*) and *Pde5a ^LoxP/LoxP^*; *Cre^Ert2^*(*blue*) fed with HFD or normal chow (NC) over 12 weeks (n= 4 each group). Data are expressed as mean ± SEM. Statistical analysis was performed applying a three-way ANOVA test.

To explore this further, we crossed *Pde5a^LoxP/LoxP^* mice with transgenic mice expressing Cre recombinase driven by the adipocyte protein-2 (*aP2/Fabp4*) promoter (*aP2^Cre^*), that is activated earlier in developing adipocytes (*Pde5a^Fabp4-cKO^*)^40^ (**Figure 8E and S4C**). *Pde5a^Fabp4-cKO^* mice displayed a greater reduction in VAT *Pde5a* expression levels (>40%) than *Pde5a*^Adpn-cKO^, and an even greater reduction (85%) in *Pde3b* levels (**Figure 8F-G**), once again confirming the interaction between the two enzymes. Unexpectedly, no differences in fat mass were observed between *Fabp4^Cre^* positive mice and control littermates (**Figure S4D),** even after 12 weeks of HFD feeding (**Figure 8H**). This suggests that protective metabolic effect of PDE5A deletion may require complete suppression of PDE5A activity, or that as specific developmental window is critical for these changes.

### Postnatal deletion of Pde5a fails to protect from HFD

To test the latter hypothesis, we crossed *Pde5a^LoxP/LoxP^* mice with *Rosa26^CreERT^* mice, allowing for tamoxifen-induced, ubiquitous postnatal deletion. Although tamoxifen markedly reduced *Pde5a* in all tissues, including VAT (**Figure 8I-K)**, where a concomitant reduction in *Pde3b* also occurred (**Figure 8K)**, these mice were not protected from HFD-induced metabolic consequences (**Figure 8L**). This suggests that the metabolic reprogramming that confers protection against HFD, must occur early during embryonic development.

To identify the critical developmental window for the observed metabolic phenotype, we isolated embryonic fibroblasts from wild-type and *Pde5a* ko embryos at 13.5 post-conception (dpc). These cells serve as a robust source of multipotent cells with multilineage potential and, under appropriate culture conditions, can effectively differentiate into adipogenic, chondrogenic, and osteogenic lineages^41^. MEFs from *Pde5a* ko mice display a lower proliferation rate compared to wild-type, but increased differentiation capacity into adipocytes, particularly at early stages adipogenic differentiation **(Figure S4E-F)**. qPCR analysis revealed upregulation of *Ppary* (peroxisome proliferation-activated receptor gamma), a master regulator of adipogenesis suggesting that PDE5A plays a role in the fine-tuning of embryonic adipogenesis (**Figure S4G**).

### Gene network analysis confirms metabolic reprogramming

RNA sequencing (RNA-seq) of VAT and liver from *Pde5a* ko mice confirmed significant transcriptional changes. In VAT 643 genes were differentially expressed, with upregulation of pathways related to thermogenesis. The Gene Ontology (GO) analysis indicated that *Pde5a* ko VAT showed significant upregulation of GO terms associated with thermogenesis, cAMP-signaling, AMPK signaling, D-amino acid metabolism, carbohydrate metabolism, and pyruvate metabolism, alongside downregulation of glycolysis/gluconeogenesis and lipid biosynthesis (**Figure S5A-C**). Notably, we found several AMPK signaling related-genes (*Map2k2, Rps6kb1, Atf4, Ngf, Hcar1* and *Arrb1*) significantly upregulated in *Pde5a* deficient VAT (**Figure S5A**).

Compared to wild-type, analysis of BAT from *Pde5a^-/-^* mice revealed 538 differentially expressed genes, with 382 genes upregulated and 156 downregulated. GO analysis revealed an increase in genes linked to improved carbohydrate metabolism, lipid metabolism, glutathione metabolism, glycolysis/gluconeogenesis, and glycine, serine and threonine metabolism, as well as adipocytokine and insulin signaling. Conversely, gene associated to NF-kappa B signaling pathway and processes related to the degradation of valine, leucine and isoleucine were downregulated in BAT from *Pde5a* ko mice (**Figure S5D-F**). Like in white adipose tissue, many genes related to AMPK signaling (*Cacna1s, Hcn2, Creb3l3, Pik3r5, Cacna1d, Cd14, Adra2b, Plcb4* and *Cacna2d1*) were significantly upregulated in brown adipose tissue from *Pde5a^-/-^* mice (**Figure S5D**) suggesting a constitutive AMPK activation.

Finally, RNA-se analysis of liver sample from *Pde5a*^-/-^ identified 477 differentially expressed genes, with 189 genes upregulated and 288 downregulated compared to wild-type mice. GO analysis revealed that AMPK and insulin signaling pathways, as well as energy metabolism and oxidative phosphorylation processes were strongly induced. Conversely pathways related to insulin resistance, steroid hormone biosynthesis, linoleic acid metabolism and amino acids metabolism signaling were repressed (**Figure S5G-I**).

In summary, both NMR and transcriptomic analyses confirm *Pde5a*^-/-^ mice undergo substantial metabolomic reprogramming in both adipose tissue and live, highlighting a global activation of AMPK signaling and a shift in metabolic pathways toward energy expenditure and away from lipid biosynthesis.

## DISCUSSION

The global prevalence of obesity is rising at an alarming rate, highlighting the urgent need to understand adipose tissue biology to develop effective therapies.^2,42^. Increasing evidence suggests that activating BAT and promoting WAT browning are promising strategies to combat obesity^2,18,43^. In this study, we unveil a previously unknown role for PDE5A in energy homeostasis using four distinct *Pde5a* knockout mouse models. The first unexpected finding was that the global constitutive knockout of *Pde5a* does not adversely affect development and reproduction; the mice behave normally and exhibit an unremarkable macroscopic phenotype. However, microscopic analysis reveals significant morphological changes in both white and brown adipose tissue, which turn out to provide robust protection against metabolic challenges.

Adipocytes from *Pde5a^-/-^* mice are significantly smaller and more numerous than wild-type, due to trans-differentiation of WAT into a more metabolically active adipose tissue, typically triggered by cold exposure or prolonged β-AR stimulation. This trans-differentiation occurs spontaneously in our mouse model, with adipocytes displaying features of *‘beige’* adipocytes”, such as increased expression of *Ucp1*, mitochondrial activity, and thermogenic potential^43,19^. Under adrenergic or cold stimuli, *Pde5a* knockout mice show reduced WAT mass and increased surface temperature, confirming thermogenic activation.

The browning of adipocytes^44^ confers *Pde5a^-/-^* mice resistance to liver steatosis, weight gain, and insulin resistance induced by aging or HFD. Metabolomic analysis revealed lower circulating and hepatic lipid levels, both basally and post-challenge. corroborated by MRI imaging showing reduced liver fat and enhanced water diffusion. Decreased pro-inflammatory cytokines and a favorable adipokine profile, characterized by increased adiponectin and reduced leptin expression, contribute to a lower leptin/adiponectin ratio (LAR), a marker inversely related to liver steatosis and diabetes risk^45,46^.

Despite unchanged food intake and physical activity, *Pde5a^-/-^*mice exhibit enhanced energy dissipation, favoring heat production over fat accumulation. This mechanism is minimal under basal conditions but activated during metabolic challenges, explaining the absence of body or fat mass differences in resting conditions. During fasting, approximately 80% of endogenous glucose production comes from liver releases. After a glucose load, insulin suppresses hepatic glucose production and release^47^. Using NMR spectroscopy, we documented an increase in Glucose-1P in *Pde5a^-/-^* mice, suggesting their liver prioritize glucose for glycogen synthesis. Lower lactate and alanine levels, coupled with reduced ^13^C labeling in glycolytic intermediates and glycerol moieties, further support this shift towards glycogen synthesis over glucose usage. The reduction of liver and circulating triglycerides suggest that *Pde5a* deficient mice rely on alternative energy sources, particularly lipids, under resting conditions.

cGMP is emerging as a key regulator of glucose and lipid metabolism^8,9^. While basal cGMP levels in *Pde5a^-/-^* mice are challenging to detect, increased PKG-dependent phosphorylation of VASP^Ser2^^39^ suggests enhanced cGMP signaling. This affects glucose metabolism by modulating acetyl-CoA carboxylase (ACC), which converts acetyl-CoA into malonyl-CoA, a precursor for de novo lipogenesis^48^; whereas, liver ketone bodies synthesis relies on Acetyl-CoA primarily derived from fatty acid oxidation^49^. Consequently, reduced ketone body concentrations in *Pde5a^-/-^* mice indicate a shift of acetyl-CoA flux towards the TCA cycle rather than hepatic ketogenesis. RNA sequencing confirms upregulation of AMPK signaling in adipose tissue and liver, aligning with its role as a master regulator of lipid metabolism through ACC1 inhibition^50^. Since ACC1 loss promotes browning^51^, AMPK activation in *Pde5a^-/-^* mice likely contributes to their thermogenic phenotype. Although we did not observe significant differences in ACC transcripts in *Pde5a^-/-^* mice, supporting evidence also derives from the elevated bile acids (BA) levels found in *Pde5a^-/-^* mice. BA are known to repress gluconeogenesis and enhance energy expenditure^52,53^, and via the TGR5 receptor stimulate liver AMPK^53^, endothelial NO-production and cGMP accumulation^54^.

Since intracellular cAMP in adipose tissue is primarily managed by *Pde3b*^30,31^, we examined its expression and found a significant downregulation, resulting in elevated cAMP levels. Targeted inactivation of *Pde3b* has been shown to enhance cAMP/PKA and AMPK signaling pathways, promoting adipose tissue browning^30,31^. The similarity between the *Pde5a^-/-^*and *Pde3b^-/-^* models suggests that the metabolic advantages of *Pde5a* deletion arise from cGMP-cAMP signaling convergence. Studies have demonstrated that increasing cAMP and cGMP levels reduces adipogenesis, upregulates UCP1, and prevents obesity under HFD^55^. *Pde10a* inhibition induces browning of human white adipocytes^56,57^, and *Pde4b^-/-^*mice appear leaner, with lower fat pad weights, smaller adipocytes, and decreased serum leptin levels on both chow and HFD^58^.

We uncovered a previously unrecognized interdependency between PDE5 and PDE3 in adipose tissue, demonstrating that: (1) *Pde5a* ablation downregulates *Pde3b* via the transcriptional repressor ICER; (2) PDE5A stabilizes PDE3B protein, preventing its degradation; and, reciprocally, (3) *Pde3b* silencing suppresses *Pde5a* transcription. However, *Pde5a^-/-^* and *Pde3b^-/-^* ko models are not entirely overlapping. First, the *Pde3b^-/-^*ko induced the beige phenotype only in SvJ129 mice, whereas in a C57BL/6 background, a β3-AR agonist was required to induce WAT browning ^59^. Second, the *Pde3b^-/-^* ko exhibited signs of insulin resistance, likely due to PGC-1α activation in the liver, whereas PGC-1α was unaffected or even reduced in *Pde5a^-/-^* BAT. Finally, *Pde3b^-/-^* models exhibited increased mitochondrial biogenesis with enlarged mitochondria, while in *Pde5a^-/-^,* mitochondria remained unaffected.

Despite heightened cAMP levels, *Pde5a^-/-^* mice do not exhibit increased lipolysis, suggesting an adaptive response. Indeed, forskolin stimulation fails to induce lipolysis in *Pde5a^-/-^* adipocytes, implicating impaired adenylate cyclase activity. *Adcy3*, the primary AC isoform in adipose tissue, is significantly downregulated in *Pde5a^-/-^* mice, likely as a compensatory mechanism to counteract excessive cAMP signaling and maintain metabolic balance. This explains why *Pde5a* deletion does not induce weight loss under normal feeding conditions.

The role of cGMP in metabolic homeostasis is further supported by *Pde9a* deletion^16^, which enhances BAT respiration, increases resting energy expenditure, and mitigates weight gain under HFD. While the metabolic benefits of *Pde9a^-/-^* and *Pde5a^-/-^* mice share similarities, the impact of *Pde9a* deletion on the cAMP-PKA axis remains unexplored.

*Pde5a* is widely distributed across tissues^4^, suggesting that the protective metabolic are not solely adipocyte-dependent. Indeed, adipocyte-selective *Pde5a* deletion (achieved using *Adipoq^Cre^* and *Fabp4^Cre^* mice) does not replicate the global knockout phenotype, implicating a role for other cell populations, possibly within the vascular stromal fraction of adipose tissue. Furthermore, inducible global *Pde5a* deletion (using *Rosa26^CreErt2^*) in adulthood fails to confer full HFD protection, suggesting that the observed reprogramming originates during early development. Alternatively, the “metabolic reprogramming” originates from a coordinated interactions among various tissues and organs. This finding reconciles discrepancies in several clinical trials^10,14^, where sildenafil improves metabolic parameters but does not significantly reduce fat mass. A recent trial administering tadalafil to diabetic patients for 12 weeks showed multiple metabolic improvements without affecting body weight^60^.

### Conclusions

This study demonstrates that mice lacking *Pde5a* are resistant to aging and HFD-induced obesity and associated liver damage, primarily due to white-to-beige conversion of adipocytes, enhanced thermogenic potential, and improved whole-body glucose and lipid homeostasis. These protective effects are mediated through a previously unrecognized interaction between *Pde5a* and *Pde3b*, which activates cAMP-PKA signaling, driving metabolic reprogramming. Our findings position PDE5A as a potential therapeutic target to enhance adipose thermogenesis, offering a promising adjuvant strategy for combating metabolic disorders.

## MATERIAL AND METHODS

### Mice generation

*Pde5a* knockout (*Pde5a^-/-^*) mice were generated using CrispR-Cas9 technology according to the strategy depicted in **Figure S1A**. sgRNAs for targeting murine *Pde5a* exon 2 were designed using the online tool MIT CRISPR Design (crispr.mit.edu). Guide RNAs were generated in vitro using the MEGAshortscript T7 transcription Kit (Thermo Fisher Scientific, MA, USA) and purified by using the MEGAclear Transcription Clean-Up Kit (Thermo Fisher Scientific, MA, USA). sgRNA (50 ng/μl), ssDNA donor oligo (100 ng/μl) and Cas9 mRNA (100 ng/μl; TriLink Biotechnologies, CA, USA) were microinjected into one-cell-stage zygotes obtained from C57BL/6N mice for the generation of Pde5 knockout mice. C57BL/6N-B6D2F1/J chimeric mice were backcrossed to obtain a pure C57BL/6N background. Three founder lines were obtained and used for characterization. All procedures were conducted in compliance with the Guidelines for Animal Care and Treatment of the European Union and Italian Law (D.Lgs 2010/63EU) and were approved by the Sapienza University’s Animal Research Ethics Committee, the Italian Ministry of Health (Authorization no. 145/2017-PR) and the Committee of Animal Care and Use of the National Cancer Institute in Frederick, Maryland.

*Pde5a* conditional knock out mice (*Pde5a^LoxP/LoxP^*) were generated using homologous recombination and BAC (bacterial artificial chromosome) technologies according to the strategy depicted in **Figure S1B.** BAC RP23-255K12 (Source BioScience Life Sciences), which consists of a 190 kb insert, including 100 kb of sequence upstream of the *Pde5a* start codon and 90 kb of mouse *Pde5a* gene, was used to generate targeting vector. Bacteria containing BAC RP23-255K12 were electroporated with mini-λ prophage DNA containing the essential components for homologous recombination. Targeting vector was generated by homologous recombination inserting the neomycin cassette, obtained from the pLTM260 vector (a generous gift of Dr. S.K. Sharan, NCI-FCRF, Frederick, MD) flanked by *LoxP* and *FRT* sites and two homology regions of the *Pde5a* gene at intron 2, into the BAC. A third *LoxP* site was introduced at intron 1. Kanamycin resistant recombinants were used for getting the BAC sequence (from intron 1 to intron 2) and the Neo cassette into the retrieval pDTa8 vector (a generous gift of Dr. S.K. Sharan, NCI-FCRF, Frederick, MD) containing the DT (diphtheria toxin) gene for negative selection of electroporated ES (embryonic stem cells). Targeting vector was linearized with NotI and electroporated into 129/Sv-C57BL/6N ES cells. Positive selection of recombinant clones was achieved using 200 μg/ml G418 (Thermo Fisher Scientific, MA, USA). 384 resistant clones were obtained and screening by Southern Blotting using two external 5’ and 3’ probes (SpeI) and an internal probe (Pvu II). An ES targeted clone was injected into C57BL/6N blastocysts and finally returned to pseudo-pregnant CD1 mothers to produce germ-line chimeric pups. Neomycin cassette was removed crossing homozygous mice with Flp-C57BL/6N deleter mice (kindly provided by Prof. D. O’Carroll, EMBL Monterotondo, Italy). AdipoQ^Cre^ and Fabp4^Cre^ mice were kindly provided by Prof. M. Riminucci (Sapienza University of Rome, Italy) while Rosa26^CreERT^ were kindly provided by Prof. S. Dolci (University of Rome Tor Vergata, Italy).

Pde5^-/-^ mice genotyping was performed using the Phire Tissue Direct PCR Master Mix (Thermo Fisher Scientific, MA, USA) and following primers: *Pde5a* screen F1: 5’-GAGTCTAGGATAGGAGCACT-3’’, *Pde5a* screen F2: 5’-TTGGCAAGGAATGTGGCTA-3’ and *Pde5a* screen R: 5’-GCAGGCTTGTTATTTACTTATTTTG-3’. Pde5^LoxP/LoxP^ mice genotyping was performed using the same Master Mix and the following primers: *Pde5a* del F: 5’-CCAGGTCATGAGAATGCACA and *Pde5a* del R: 5’-TGTCTCTTGGACTGGCAGA.

All mice used for experiments were provided with food and water ad libitum and kept in a standard specific pathogen-free environment.

All procedures were conducted in compliance with the Guidelines for Animal Care and Treatment of the European Union and Italian Law (D.Lgs 2010/63EU) and were approved by the Sapienza University’s Animal Research Ethics Committee, the Italian Ministry of Health (Authorization no. 165/2016-PR) and the Committee of Animal Care and Use of the National Cancer Institute in Frederick, Maryland.

### High-fat diet feeding protocol

A mouse obesity model was established through chronic feeding with a HFD (60% kcal from fat, D12492, Research Diets, NJ, USA) for 12 weeks starting at the age of 8 weeks. Normal chow diet-fed mice were maintained on diet with 10% kcal from fat (D12450J, Research Diets, NJ, USA). Food and water were provided ad libitum. Mice were housed and kept on a 12h light/dark cycle at 23□±□1□°C and weighed before HFD induction and once a week for all the duration of HFD feeding protocol. Food and water intake was determined weekly for all the duration of the experiment. Food Efficiency Ratio (FER) was calculated as: FER = 100*(weight gain/food intake) as previously described ^61^.

### *In vivo* thermogenesis induction

For cold-induced thermogenesis 8-week-old male mice (n=5/group) were housed at a constant room temperature of 4□°C for 4 hours according to^43^.

For chemically induced thermogenesis 8-week-old male mice (n=6/group) were injected via *ip* with 1 mg/kg β3 adrenoceptor agonist CL316,243 (Sigma-Aldrich, MA, USA) or saline for 10 days as previously described^62^. Mice were weighted before and at the end of the treatment. Mice were housed at a constant room temperature of 23□±□1□°C with food and water given ad libitum and were fed a standard diet that contains 62% kcal from carbohydrate, 25% kcal from protein, and 13% kcal from fat (Inotiv, IL, USA).

### IR thermography

Mice were housed in individual cages in an air-conditioned room maintained at a temperature of 22–24°C. The infrared camera (Teledyne FLIR, Boston, USA) was mounted on a tripod positioned 40 cm above a clean and dry cage without the lid. The temperature data were recorded on a memory card in radiometric CSQ format. The infrared videos were transferred and analyzed by using thermal camera software (FLIR Tools+ version 5.3.15320.1002, FLIR Systems). The absolute values of radiative temperature in the interscapular skin, representing the interscapular BAT depots, and the differences between temperatures of the interscapular back skin and the lumbar back skin (as the reference temperature) have been considered, as described elsewhere^63^. For cold exposure experiments, ANCOVA test was performed comparing post-exposure temperature values and accounting for baseline temperature values as covariate.

### Histological analyses

Tissues were fixed in 10% neutral buffered Formalin (Sigma-Aldrich, MA, USA), dehydrated with increased grade alcohols and embedded in Paraffin (Bio Optica, Italy). 5μm sections obtained with the HM355S Microtome (Thermo Fisher Scientific, MA, USA) were de-waxed, re-hydrated, and finally stained with Hematoxylin and Eosin (Sigma-Aldrich, MA, USA). Slides were mounted and images were acquired using a Zeiss Axiovert 200 inverted microscope (Carl Zeiss Inc., NY, USA) equipped with a AxioCam 503 Color (Carl Zeiss Inc., NY, USA). Fat coefficient was expressed in percentage and calculated by relating the weight of each AT depot (visceral adipose tissue-VAT refers to epididymal fat, subcutaneous adipose tissue - SAT refers to inguinal fat, brown adipose tissue - BAT refers to interscapular brown adipose tissue) to body weight. Cross-sectional area estimates (100 cells) belonging to three independent experiments were used for adipocyte area determination.

### Immunohistochemistry

Immunohistochemistry was performed as previously described^64^. Briefly, formalin-fixed paraffin-embedded (FFPE) sections obtained with the HM355S Microtome (Thermo Fisher Scientific, Waltham, MA, USA), were de-waxed, re-hydrated and finally processed for IHC using the EnVision®+ Dual Link System-HRP (DAB+) (DAKO/Agilent, Santa Clara, CA, USA) according to manufacturer’s instructions. Washes were performed three times with PBS + 0,05% v/v Tween20. Antigen retrieval was performed by microwaving sections in 10 mM Sodium Citrate pH 6.0 + 0,05% Tween20 v/v for 10 min. Sections were incubated overnight at 4°C with UCP1 antibody (see antibody list) diluted in Bond Primary Antibody Diluent (Leica, Wetzlar, Germany). Before mounting, slides were counterstained with Hematoxylin (Sigma Aldrich, Saint Louis, MO, USA). Images were acquired by Zeiss Axiovert 200 inverted microscope using ZEN imaging software (Carl Zeiss., Oberkochen, Germany).

### Western Blotting

Tissues were lysed in RIPA Buffer (Sigma-Aldrich, MA, USA) plus protease and phosphatase inhibitor cocktail (Sigma-Aldrich, MA, USA). Total protein amount was determined using the BCA protein assay kit (Thermo Fisher Scientific, MA, USA). Laemmli sample buffer (BioRad Laboratories, CA, USA) was added to lysates and samples were boiled for 5 min at 95 °C. Denatured samples were electrophoresed in polyacrylamide gels and transferred onto PVDF membranes (Amersham, NY, USA). Primary antibodies are listed in **Antibodies List** and were incubated overnight at 4 °C and secondary antibodies were incubated for 1□hour at room temperature. Signals were detected with horseradish peroxidase (HRP)-conjugated secondary antibodies and enhanced chemiluminescence (Thermo Fisher Scientific, MA, USA). Chemiluminescent images of immunodetected bands were recorded with the Syngene G-box system (Syngene Bioimaging, MD, USA), and immunoblot intensities were quantitatively analyzed using ImageJ Software (NIH, Bethesda, MD, USA). Results represent the means of at least three independent experiments, and were normalized to the amount of housekeeping proteins.

### Antibodies List

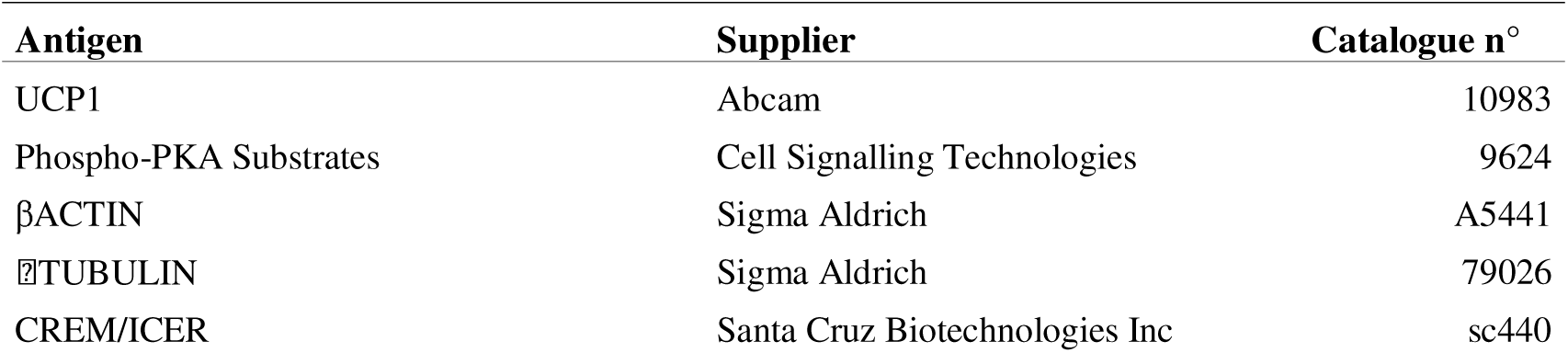
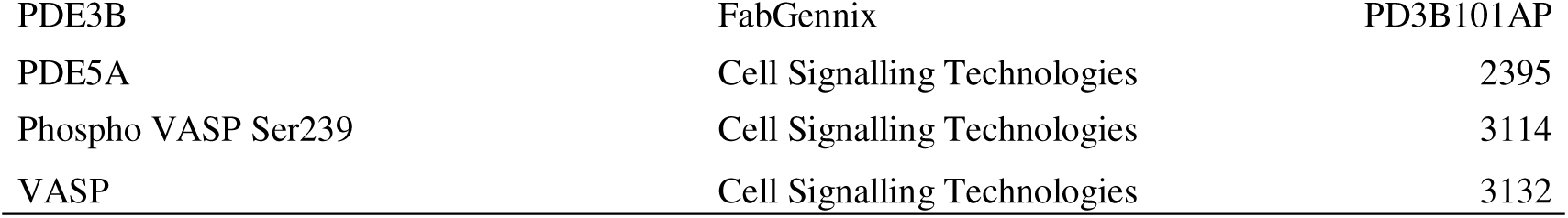

### PDE activity and cAMP assay

PDE activity, expressed as μmoles cGMP hydrolyzed/min/mg of enzyme, was measured at 30°C with the two-step method described by Thompson and Appleman, using ^[3]^H cGMP (Perkin Elmer, MA, USA) as previously described^4^. Mouse embryonic fibroblasts at passage two were homogenized using a glass homogenizer (15 strokes, 4°C) in 20 mM Tris-HCl buffer pH 7.2 containing 0,2 mM EGTA, 5 mM β-mercaptoethanol, 2% v/v protease inhibitor cocktail (Sigma– Aldrich, MA, USA), 1 mM PMSF, 5 mM MgCl_2_ and 0,1% v/v Triton X-100. The homogenates were centrifuged at 14,000g for 30 min at 4°C and pellets were re-suspended in the homogenization buffer and centrifuged at 14,000g for 30 min at 4°C. Extracts were incubated in 60 mM HEPES pH 7.2 assay buffer containing 0,1 mM EGTA, 5 mM MgCl_2_, 0,5 mg/ml bovine serum albumin, 30 μg/ml soybean trypsin inhibitor, in a final volume of 0,3 ml. The reaction was started by adding tritiated substrate at a final concentration of 1 μM and stopped by adding 0,1 M HCl. The specific activity was quantified at the 10% limit of the total substrate hydrolyzed. Sildenafil citrate (Pharmaceutical PolPharma, Poland) was used as previously described^4^. cAMP levels were assayed in adipose tissue extracts using the Direct cAMP ELISA kit (Enzo Life Sciences Inc., NY, USA) according to manufactures’ instructions for acetylated form. Determination of the protein amount in the cyclic nucleotide samples was evaluated in neutralized acid homogenate using the BCA protein assay kit (Thermo Fisher Scientific, MA, USA).

### Metabolic profiling

Cellular oxygen consumption rate (OCR) and extracellular acidification rate (ECAR) were detected using XF Cell Mito Stress Test (Agilent Technologies, CA, USA) measured by the extracellular flux analyzer XFe96 (Seahorse Bioscience, TX, USA) in the HypACB facility at Sapienza University of Rome. Primary adipocytes were seeded at a density of 7x10^3^ cells/well and cultured on XFe culture 96-wells miniplates for 24 hours. The sensor cartridge for XFe analyzer was hydrated in a 37□°C non-CO_2_ incubator a day before the experiment. According to the manufacturer instructions, stressors concentrations were optimized and added as follows: 1□μM oligomycin as complex V inhibitor, 1□μM FCCP as uncoupler agent and 0,5□μM rotenone/antimycin A (inhibitors of complex I and III). During sensor calibration, cells were incubated in a 37□°C non-CO_2_ incubator in 180 μl assay medium (XF base medium supplemented with 10□mM glucose, 10□mM pyruvate and 2□mM L-glutamine at pH 7.4, was used to wash the cells and replace the growth medium). OCR and ECAR were normalized for total protein/well/10^4^ cells. Each sample/treatment was analyzed in 8 wells for experiment. Two independent experiments were carried out. Statistical analysis has been performed on each experiment by using one way ANOVA followed by Bonferroni post-hoc comparison test. The differences were considered statistically significant when p<0.05.

### RNA sequencing

RNAseq analysis was performed on total RNA extracted from adipose tissue (VAT, SAT and BAT) and liver of 8-week-old male mice (n=3 *Pde5a^+/+^* and n=3 *Pde5a^-/-^*) fed with standard chow as well as HFD fed mice (n=3 *Pde5a^+/+^* HFD and n=3 *Pde5a^-/-^* HFD). Total RNA was isolated using RNeasy Lipid Tissue mini kit (Qiagen, Germany) according to manufacturer’s instruction. RNA was treated with DNAse Qiagen, Germany) and RNA samples were quantified and quality tested by Agilent 2100 Bioanalyzer RNA assay (Agilent technologies, CA, USA) or by Caliper LabChip GX (PerkinElmer, MA, USA). Universal Plus mRNA-Seq kit (Tecan Genomics, CA, USA) has been used for library preparation (fr-secondstrand) following the manufacturer’s instructions. Final libraries were checked with both Qubit 2.0 Fluorometer (Invitrogen, CA, USA) and Agilent Bioanalyzer DNA assay or by Caliper LabChip GX (PerkinElmer, MA, USA). Libraries were then prepared for sequencing and sequenced on paired-end 150 bp mode on NovaSeq 6000 (Illumina, San Diego, CA). Base calling, demultiplexing and adapter masking were performed using Illumina BCL Convert v3.9.31. The adapter sequences were masked during the demultiplex with BCL-Convert. Trimming was performed using ERNE2 software. Reads were aligned on the reference genome (*mm10*) with STAR3. Assembling and quantitation of full-length transcripts representing multiple spliced variants for each gene locus was performed using Stringtie 4. Statistical significance was analyzed using the student’s t-test for parametric data and a combined t-test/ ANOVA test with U Mann-Whitney correction for nonparametric data following a non-homoscedastic distribution. The Shapiro-Wilk test was employed to assess data normality prior to the t-test, followed by the Brown Forsythe test for checking homoscedasticity condition for each variable, with a significance value of 0.05. If these conditions were not met, the Wilcoxon rank sum test was performed. Spearman’s Clustergram was also done for checking pattern with significant variables and samples. All differences were considered statistically significant with p<0.05. Analyses were performed using MATLAB® R2023a (MathWorks, Natick, Massachusetts, USA) with the Statistics and Machine Learning Toolbox and Bioinformatics package.

### Body composition analysis

Abdominal MRI analysis was performed on 12-month-old male mice (n=7 *Pde5a^+/+^* and n=7 *Pde5a^-/-^*) using the BioSpec 94/20 USR scanner 9.4T (Bruker, MA, USA). Following cervical dislocation, each mouse was placed in sternal recumbency on MRI bed for post-mortem examination. The protocol consisted in a T2-weighted sequence with fat-water separation (voxel size 0.13 x 0.13 x 0.5 mm^3^; 24 slices) and a Diffusion weighted sequence (voxel size: 0.47 x 0.47 x 0.5 mm^3^; 10 slices; b-values; 0, 150, 500, 800) both acquired on a transverse plane. The T2-weighted sequence with fat-water separation yielded two images per slice: water only and fat only images. In order to quantify the fat accumulation in abdominal organs, the two image volumes were spatially aligned with an affine registration implemented with Elastix^65^, to correct for the displacement caused by the fat-water shift. They were then processed in order to enhance the Signal-to-Noise Ratio (SNR) with a revised version of the Multi-spectral Non-Local Means (MNLM) approach was used^66^. Finally, a voxel-by-voxel Fat Fraction (FF) parameter map was calculated by means of an in-house software, according to the equation FF(%) = F/(F + W), where *F* was the fat signal and *W* the water signal^67^. From the FF map, through a signal thresholding and morphological operation, an automated segmentation of the subcutaneous and visceral fat was obtained. From this binary mask, a total volume was automatically computed. In order to evaluate the water microscopic mobility within the liver parenchyma, the images corresponding to the 4 b-values of the abdominal DWI were first denoised by the MNLM approach, then they were used to compute the Apparent Diffusion Coefficient (ADC) map.

Image analysis was performed using in-house software developed in MATLAB®. GraphPad Prism 9 was employed for the statistical analysis, using a t-test for the group comparison. The differences were considered statistically significant when p<0.05.

### Gene expression analysis

Total RNA was isolated from adipose tissue biopsies using RNeasy Lipid Tissue mini kit (Qiagen, Germany) according to manufacturer’s instruction. RNA was treated with DNAse Qiagen, Germany) and reverse transcribed with random hexamer primers using Maxima cDNA synthesis kit (Thermo Fisher Scientific, MA, USA). Real time PCR reaction was carried out in triplicate for each gene and sample by using PowerUP SYBR Green Master Mix (Thermo Fisher Scientific, MA, USA). The PCR reaction was carried out using the QuantStudio 7 System (Thermo Fisher Scientific, MA, USA). Primers pairs are listed in **RTqPCR primers list**. For quantification analysis, the comparative threshold cycle (Ct) method was used. The Ct values of each gene were normalized to the Ct value of *Hprt1*. The gene expression levels were evaluated by the fold change using the equation 2^−ddCt^.

### RTqPCR primers list

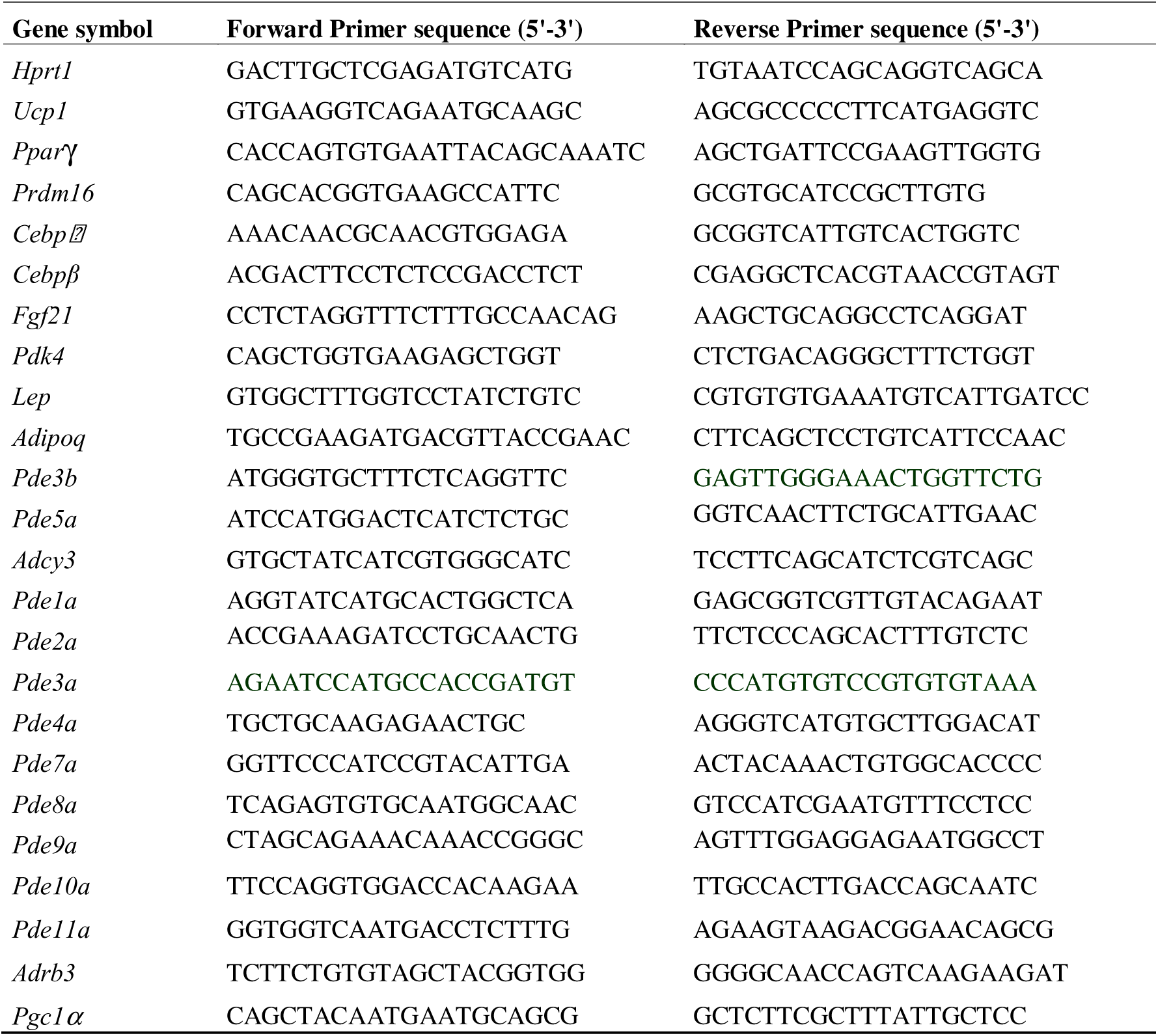

### Intraperitoneal Glucose Tolerance Test (IGTT)

Glucose tolerance test was performed on 8-week-old male mice (n=5/group) as previously described^68^. Mice were fasted overnight and 2 g/kg of Glucose (Sigma–Aldrich, MA, USA) dissolved in saline were delivered by *i.p.* injection. Blood glucose levels were monitored from the tail-tip using the Multicare-In Strips System (BSI Diagnostics, Italy) in the basal state and 15, 30, 60, 120 and 180 min following glucose administration.

### Intraperitoneal Insulin Tolerance Test (IITT)

Insulin tolerance test was performed on 8 weeks old male mice (n=5/group) as previously described^68^. Mice were fasted overnight and 0,5 U/kg of Insulin (Sigma–Aldrich, MA, USA) dissolved in saline were delivered by *i.p.* injection. Blood glucose levels were monitored from the tail-tip using the Multicare-In Strips System (BSI Diagnostics, Italy) in the basal state and 15, 30, 45, 60 and 120 min following glucose administration.

### Mouse embryonic fibroblasts isolation

Mouse embryonic fibroblasts (MEFs) were isolated from 13.5 day post coitum (dpc) *Pde5a* ko and wild-type embryos as previously described^41^. After the cells had reached confluency, medium was replaced with adipogenic medium: DMEM F12 (Life Technologies, CA, USA), 10% Fetal Calf Serum (FCS, Life Technologies, CA, USA), 393□ng/mL dexamethasone (Sigma Aldrich, MA, USA), 50□μg/mL ascorbic acid-2-phosphate (Sigma Aldrich, MA, USA), penicillin/streptomycin, 111,1□μg/mL 3-isobutyl-1-methylxanthine (IBMX, Sigma Aldrich, MA, USA), 1□μg/mL insulin (Sigma Aldrich, MA, USA), and 35,8□μg/mL indomethacin (Sigma Aldrich, MA, USA). Cells were cultured for up to 14 days until differentiation was reached.

### Cell viability assay

Cell viability was determined on MEFs using the MTT Assay (Sigma Aldrich, MA, USA) according to manufacturer’s instructions. 5x10^3^ cells were seeded onto a 96 wells cell culture multi well plate and assay was conducted for up to 72h. Absorbance at 570 nm was determined spectrophotometrically using a D3 Plate Reader (DAS, Italy). At least six replicates were made for each experimental group.

### Adipogenic differentiation

Mouse embryonic fibroblasts were seeded at a density of 3x10^3^ cells per cm^2^ in culture medium - DMEM high glucose supplemented with 10% Fetal Bovine Serum (FBS, Life Technologies, CA, USA), L-Glutamine and penicillin/streptomycin (Life Technologies, CA, USA) and cultured until 70% confluence was reached. Cells were then cultured for 3 days in differentiation medium: culture medium supplemented with 1□µM Dexamethasone, 10□µg/mL Insulin and 0,5□mM 3-isobutyl-1-methylxanthine (IBMX). After 3 days medium was replaced with culture medium supplemented with 10□µg/mL Insulin and cultured for 3 days. Medium was finally replaced and cells were maintained in culture medium for 2 days. Differentiation was measured by Oil Red O (ORO) staining. Cells were fixed with 10% neutral buffered Formalin (Sigma-Aldrich, MA, USA) and fixed cells were incubated with ORO (2□mg/mL, Sigma-Aldrich, MA, USA) for 30□min. After observation with a Zeiss Axiovert 200 inverted microscope (Carl Zeiss Inc., NY, USA) equipped with a AxioCam 503 Color (Carl Zeiss Inc., NY, USA), ORO was solubilized with isopropanol and quantified on a D3 Plate Reader (DAS, Italy) at a wave length of 510□nm.

### Lipolysis

Lipolysis was determined by measuring free fatty acids (FFA) and glycerol release from gonadal fat explants and serum obtained from 2-month-old male mice using Free Fatty acid Quantification Kit and Glycerol Assay Kit (Sigma-Aldrich, MA, USA). For basal lipolysis experiment, 10 mg VAT was incubated in 200 μl of 1% v/v Chloroform/Triton X100 and homogenized using the Precellys Evolution System (Bertin Technologies, Montigny-le-Bretonneux, France). Samples were centrifuged and organic phase was air dried. Lipids were dissolved in assay buffer and FFA and glycerol release was determined according to manufacturer’s instructions. For stimulated lipolysis experiments, adipose tissue was cutted into 20 mg pieces and pre-incubated at 37 °C, 5% CO_2_, and 95% humidified atmosphere for 60 min in 200 μl DMEM containing 1g Glucose, 2% BSA (fatty acid-free) and 10 μM forskolin/isoproterenol at 37°C. Medium was replaces for additional 60 min. Media were collected at indicated time points for the measurement of glycerol and FFA release. Total protein amount was determined using the BCA protein assay kit (Thermo Fisher Scientific, MA, USA) and FFA/Glycerol concentrations were normalized accordingly.

### Serum biochemical analyses

Serum fasting glucose was determined using the Multicare-In Strips System (BSI Diagnostics, Italy). Serum levels of Leptin, Adiponectin, Insulin, IL1α and INFγ were evaluated with commercially available enzyme-linked immunosorbent assay - ELISA (Thermo Fisher Scientific, MA, USA). The optical density was measured spectrophotometrically at a wavelength of 450 nm according to the manufacturer’s instructions. All analyses were performed in duplicate. A 4-PL standard curve was created using MyCurveFit Beta Software.

### Mitotracker staining

VAT and BAT pads were fixed in 10% neutral buffered Formalin (Sigma-Aldrich, MA, USA) overnight, dehydrated with increased grade alcohols and embedded in Paraffin (Bio Optica, Italy). 5μm sections obtained with the HM355S Microtome (Thermo Fisher Scientific, MA, USA) were de-waxed, re-hydrated, and stained with 500 nM MitoTracker Red chloromethyl-X-rosamine (CMXRos) (Thermo Fisher Scientific, MA, USA) for 10 min at room temperature. Slides were washed, mounted, and observed with a Nikon Eclipse Ti-S microscope (Nikon Corporation, Tokio, Japan).

### Fluorescence resonance energy transfer (FRET)

FRET analysis was performed on MEFs isolated from *Pde5a* ko and wild-type mice and transfected for 48h with Lipofectamine 3000 (Life Technologies, CA, USA) with a probe sensitive to Protein Kinase A (PKA) activation (AKAR3) as previously described^69^. Cells were stimulated with 1μM forskolin and 100 nM isoproterenol and imaged on a Nikon Ti50 microscope equipped with a charge-coupled device camera controlled by Metafluor software (Molecular Devices, Sunnyvale, CA). Fluorescence images were recorded by exciting the donor fluorophore at 430-455 nm and measuring emission fluorescence with two filters (cyan and yellow). Images were subjected to background subtraction, and were acquired every 20 seconds with exposure time of 200 ms. The donor/acceptor FRET ratio was calculated and normalized to the ratio value of baseline.

### Locomotory activity analysis

Locomotory activity was determined by novelty-induced exploration test on 8-week-old male mice (*Pde5a^+/+^,* n=7 and *Pde5a^-/-^*, n=7) as previously described^70^. Mice were individually placed in an experimental cage (L35cm x W25cm x H30cm) to monitor locomotory activity. The horizontal motor activity was evaluated at 10 min intervals over a 60 min test session through a computerized video tracking system (Videotrack, Viewpoint S.A., France).

### Cell culture

3T3-L1 MBX cells were purchased from ATCC (# CRL-3242) and cultured in DMEM high glucose supplemented with 10% Fetal Bovine Serum (FBS, Life Technologies, CA, USA) and penicillin/streptomycin (Life Technologies, CA, USA). For *Pde3b* silencing experiments, control non-targeting siRNA (ON-TARGETplus Non-targeting Pool) and siRNA directed against *Pde3b* (ON-TARGETplus Mouse *Pde3b* siRNA SMART Pool #18576) were purchased from Dharmacon (Dharmacon, CA, USA). Transfection was performed using the DharmaFECT 1 siRNA Transfection Reagent (Dharmacon, CA, USA) according to manufacturer’s protocol. Cells were harvested 72 hours after transfection for RNA extraction.

For *Pde5a* overexpression experiments, transfection was performed using Lipofectamine 3000 (Life Technologies, CA, USA) according to manufacturer’s instructions. Briefly 5×10^5^ cells/well were plated in 60 mm dishes and transfected with a *Pde5a*-GFP plasmid or with an empty GFP plasmid (pEGFP-c1, Clontech Laboratories, CA, USA). The generation of *Pde5a*-GFP plasmid was previously described^4^. 48 hours after transfection, cells were treated with 5 μg/mL cycloheximide (CHX, Sigma Aldrich, CA, USA) for indicated times (30 min to 240 min) and harvested to be lysed for western blotting analysis.

### NMR metabolomics

NMR analysis was performed on liver, kidney, serum, BAT and VAT from *Pde5a^-/-^* and wild-type mice both starved and 1 hour after 1mg/kg glucose administration via gavage. Samples were extracted following a modified Bligh-Dyer protocol^71^. Briefly, each organ was rapidly frozen in liquid nitrogen, weighted and immediately ground in a mortar with liquid nitrogen. Cold chloroform methanol, and water were added to frozen tissue powder at a final ratio of 2:2:1, respectively. The samples were stirred, stored at 4 °C overnight, and then centrifuged for 25 min at 4 °C at 10.000x g. The upper hydrophilic and the lower organic phases were carefully separated and dried under nitrogen flow. The hydrophilic phase was resuspended in 0,7 mL of D_2_O containing 3-(trimethylsilyl)-propionic-2,2,3,3-d_4_ acid sodium salt (TSP, 2 mM) as an internal chemical shift and concentration standard. The lower organic phase was resuspended in 0,7 mL of CDCl_3_ containing hexamethyldisiloxane (HMDSO, 2 mM) as an internal chemical shift and quantitative standard. Serum was subjected to the Bligh-Dyer procedure as the solid matrices, but without liquid nitrogen and adding water to the serum until a 1 mL volume is reached as previously described^71^. The NMR ^1^H monodimensional spectra were recorded at 25°C on a JEOL ECZR JNM spectrometer with a magnet operating at the proton frequency of 600.13 MHz. Spectra were acquired collecting 128 scansions for each sample using a calibrated 90° detection pulse length of 8.3 µs, 64k data points and a spectral width of 15 ppm. Presaturation has been employed for water signal suppression and the relaxation delay has been set to 7,723 s, in order to achieve a 15 s of total acquisition time, guaranteeing complete resonance relaxation between following scansions. Spectra have been processed by applying an exponential window function with a line broadening factor LB=0.3Hz. After applying the Fourier transformation, spectra have been manually phased and base corrected by applying the BCFR protocol. Metabolites quantitation has been carried out by comparing the integrals of specific resonances with the one of the internal standards (TSP or HDMSO for aqueous or organic fraction respectively) and normalized by the number of protons with the final data being expressed as µmol/g. For quantification only those signals that did not overlap with other resonances were chosen for integration according to the general formula:

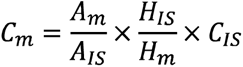

where C_m_ is the concentration of the metabolite, A_m_ is the area of the metabolite signal, H_m_ is the number of protons generating the metabolite signal, C_IS_ is Internal Standard (IS) concentration, A_IS_ is the area of the IS signal and H_IS_ is the number of protons generating the IS signal. The labelling experiments was performed using ^13^C (U-C6) glucose (CIL, Cambridge Isotope Laboratory, Massachusetts, USA). The NMR ^13^C monodimensional spectra were recorded at 25°C employing an inverse gated pulse sequence to remove NOE effect compromising the ^13^C quantitative analysis^72^. Relaxation delay was set to 7 s for all the spectra, acquiring 64 k data points in 1.38 s, to achieve the complete relaxation of ^13^C except for the ^13^C of carbonyl groups. The spectral width was set to 47,4 kHz (250 ppm) and 3000 scans were collected for each spectrum to achieve an acceptable signal-to-noise ratio. The ^13^C isotopomer analysis was encoded starting from the ^13^C NMR spectra by quantifying the ^13^C in specific positions of the carbon skeleton of the molecule on the basis of ^13^C NMR spectral multiplicity due to ^13^C-^13^C scalar coupling. The fractional ^13^C enrichment YCi is defined as the amount of ^13^C relative to the total carbon (^13^C+^12^C) present in Ci^73^, according to the following formula: YCi = ^13^Ci /(^13^Ci + ^12^Ci). To univocally identify the metabolites in the biological samples, bidimensional experiments ^1^H-^1^H Total Correlation Spectroscopy (TOCSY) and ^1^H-^13^C Heteronuclear Single Quantum Correlation (HSQC) and Heteronuclear Multiple Bond Correlation (HMBC) were performed on selected samples. TOCSY experiments were conducted with a spectral width of 9025 Hz in both dimensions, a data matrix of 8192×256 points, a mixing time of 80ms, and a relaxation delay of 2s. HSQC experiments were performed with spectral widths of 9025KHz and 37,764 kHz for the proton and carbon, respectively, a data matrix of 8192×256 points and a recycle delay of 2 s. HMBC experiments have been acquired with a spectral width of 9025 kHz and 37,764 kHz for the proton and carbon, respectively, with a data matrix of 8 K×256 points, long-range constants nJC–H of 4, 8, and 12Hz, and a recycle delay of 3s. Given the high correlation of variables resulting from a metabolomic analysis, the data obtained were first considered from a multivariate point of view. To evaluate any samples spontaneous grouping, unsupervised PCA were applied to the entire dataset after mean-centering and scaling. Besides, to highlight the role of each metabolite in distinguishing between the groups a PLS-DA model has been built, using as validation procedure the Repeated Double Cross Validation method. Accuracy, Precision, Sensitivity and Specificity were then employed as validation parameters. Significant variables for regression have been selected taking into account the Regression Coefficients, considering only variables whose Regression Coefficients sign remained consistent during cross validation procedures. All Analyses were performed using in-house routines running under the MATLAB environment (The MathWorks, Natick, MA, USA), with the Statistics and Machine Learning Toolbox and Bioinformatics package.

### Statistical analyses

Data are presented as mean ± SEM from at least three independent experiments. Data distribution was assumed to be normal but this was not formally tested.

Statistical significance was analyzed using the student’s t-test for parametric data and the ANOVA test with Tukey’s corrections for nonparametric data.

All differences were considered statistically significant when *p<0.05, **p< 0.01, ***p<0.005 and ****p<0.001.

All analyses were performed using GraphPad Prism 9 software.

## ACKNOWLEDGMENTS

This work was supported by grants from Italian Ministry of Research (grant no. P2022CE79J to FC; 2020XMLP45_004 to SD) and Sapienza University of Rome (grants no. RM123188F740EABA to F.C; AR22117A86D7E9FC to F.C; SP122184854B46B6 to F.C and RM12218167FC03E8 to

M.A.V). Sapienza University of Rome Research Infrastructures for the HypACB platform for Seahorse experiments is fully acknowledged (grant no. GA116154C8A94E3D). The authors thank the EuroBioImaging and the Multi Modal Molecular Imaging Italian Node Facility at the Institute of Biostructures and Bioimaging (CNR) Naples, Dr. Sandra Albanese and Dr. Ernesto Soscia for their support in MRI experiments and Dr. Alice Di Stefano and Dr. Tamara Antici for their contributions in IR-thermography experiments. Parts of Figure S2 and 8 were created using BioRender.com.

## AUTHOR CONTRIBUTIONS

Conceptualization: F.C. and A.M.I.; methodology and investigations: F.C., O.G., F.B., B.P., A.D.M., F.S., F.R., S.M., S.C., M.R.A., E.P., A.P. and L.T.; data analysis and interpretation: F.C., F.S., E.P., S.D., M.G., A.M. and L.T.; funding acquisition: F.C., S.D., M.A.V. and A.M.I.; project administration: F.C. and A.M.I.; draft writing: F.C. and A.M.I.; draft editing: F.C., O.G., F.B., B.P., A.D.M., F.S., F.R., S.M., S.C., M.R.A., E.P., A.P., A.F., A.L., D.G., M.G., S.D., F.N., M.S., M.R., M.M., A.M., L.T., M.A.V and A.M.I.

## DATA AVAILABILITY

The authors confirm that the data supporting the findings of this study are available within the article and/or its supplementary materials. The datasets generated during and/or analyzed during the current study are available from the corresponding author upon request.

---

**Supplementary Figure 1.**
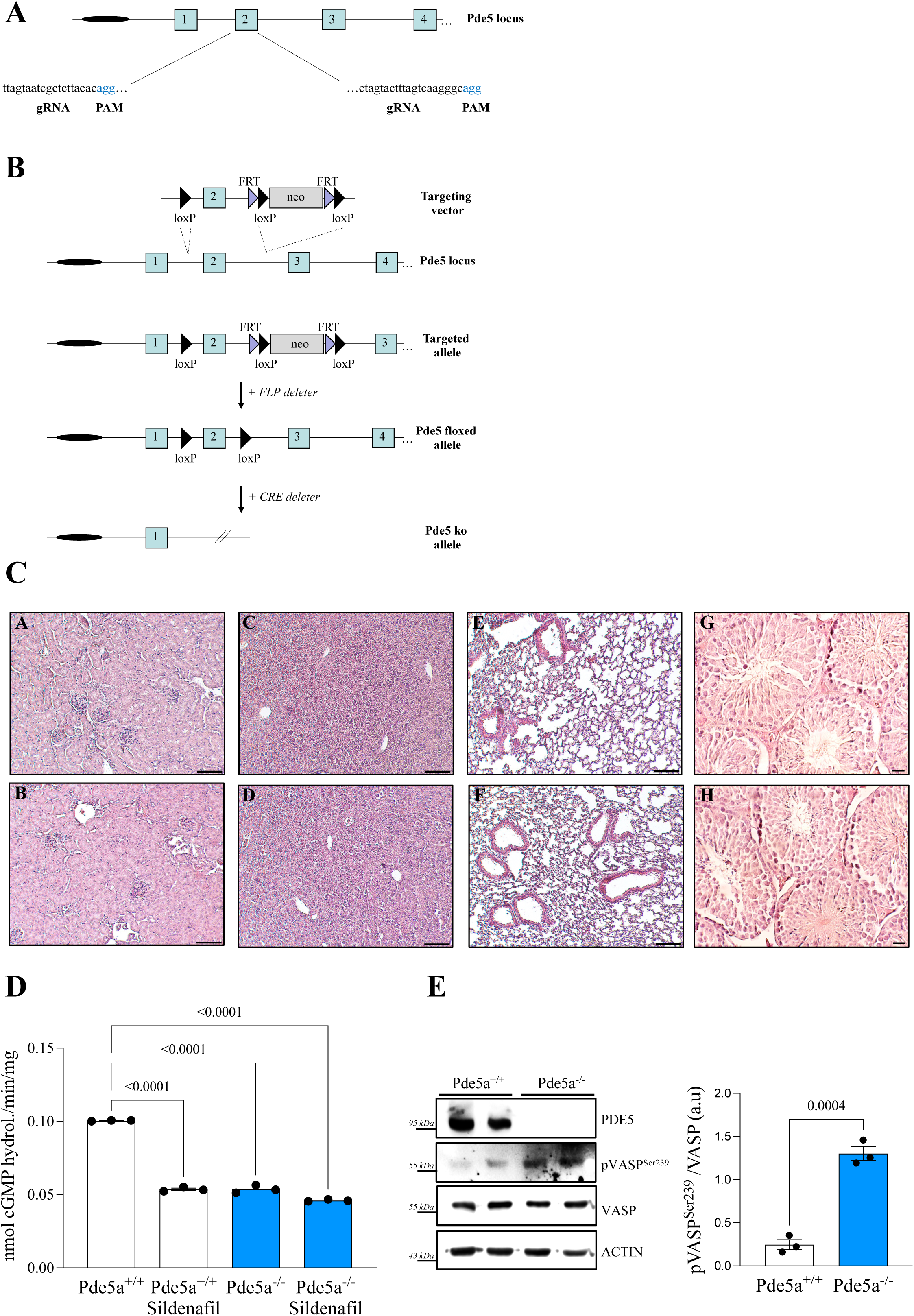
(A) Schematic representation of targeting strategy of global *Pde5a* knockout. Exons are shown in light blue. RNA guide sequences (gRNA) and PAM sequences are highlighted. (B) Schematic representation of targeting strategy of *Pde5a* conditional knockout. Exons are shown in light blue; *loxP* sites are indicated as black triangles; *Frt* sites are indicated as liliac triangles. (C) Hematoxylin and Eosin staining of kidney (a, b), liver (c, d), lung (e, f) and testis (g, h) FFPE sections obtained from *Pde5a*^+/+^ (*left panels*) and *Pde5a*^-/-^ (*right panels*) mice. Scale bar = 50 μm. (D) PDE activity, expressed as μmoles cGMP hydrolyzed/min/mg of enzyme in MEF lysates obtained from *Pde5a*^+/+^ (*white*) and *Pde5a*^-/-^ (*blue*) mice in presence or not of 1 μm PDE5i sildenafil. Data presented as mean of 3 independent experiments ± SEM. Statistical analysis was performed using two-way ANOVA test. (E) Representative immunoblots of VASP PKG-selective phosphorylation on Serine^2^^39^ and PDE5A protein levels in whole lysates obtained from visceral adipose tissue obtained from *Pde5a*^+/+^ and *Pde5a*^-/-^ (*n =3*) mice. pVASP was normalized on total VASP. Densitometric analysis is shown in the graph on the right. Data are presented as dot plots with column bars ± SEM. Statistical analysis was performed using Student t-test.

**Supplementary Figure 2.**
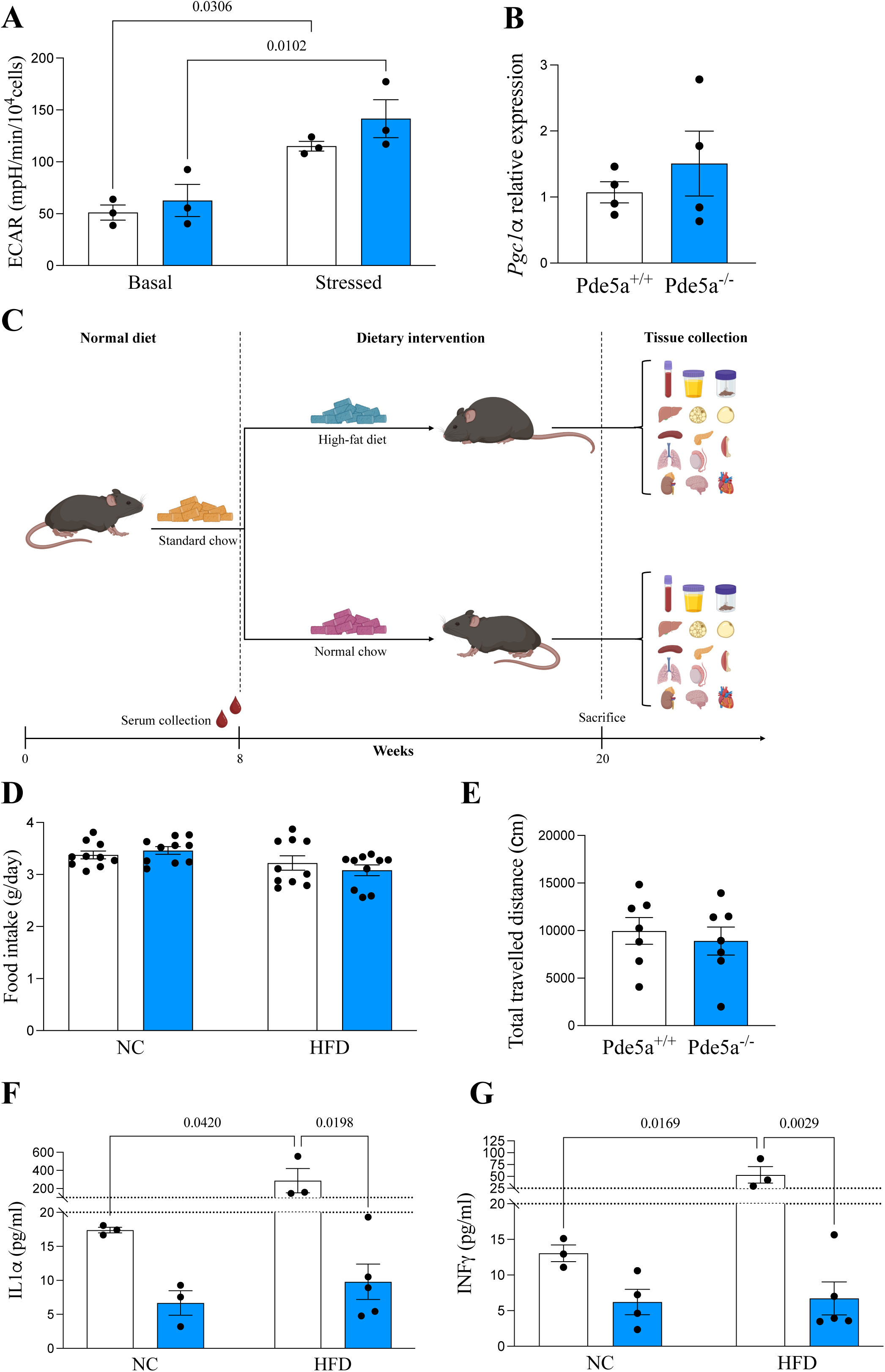
(A) Respiration and glycolysis analysis on VAT cultures from *Pde5a*^+/+^ (*n =3, white*) and *Pde5a*^-/-^ (*n =3, blue*) mice trough quantification of basal and stressed extracellular acidification rate (ECAR). Data presented as mean of 3 independent experiments ± SEM. Statistical analysis was performed using one-way ANOVA test. (B) qPCR gene expression analysis of PGC1α performed on VAT obtained from *Pde5a^+/+^* (*n =4, white*) and *Pde5a^-/-^*(*n =4, grey*) mice. *Hprt1* was used as housekeeping gene for normalization. Data are presented as dot plots with column bars ± SEM. Statistical analysis was performed using Student t-test. (C) Dietary challenge experimental protocol showing time, treatment and outcomes. (D) Measurements of total daily food intake in *Pde5a^-/-^* (*blue*) and WT (*white*) mice after 12 weeks feeding with NC (n=10) and HFD (n = 10). Data are presented as dot plots with column bars ± SEM. Statistical analysis was performed using two-way ANOVA test. (E) Habituative profile of locomotion measured as total travelled distance of *Pde5a*^-/-^ (*blue*) and WT (*white*) mice. Data are presented as dot plots with column bars ± SEM. Statistical analysis was performed using one-way ANOVA test with n= 7. (F-G) Analysis of serum levels of IL1α (F) and INFγ (G) trough ELISA on *Pde5a*^-/-^ (*blue*) and wild-type (*white*) mice after 12 weeks feeding with NC (n=3) and HFD (n = 5). Data are presented as dot plots with column bars ± SEM. Statistical analysis was performed using two-way ANOVA test.

**Supplementary Figure 3.**
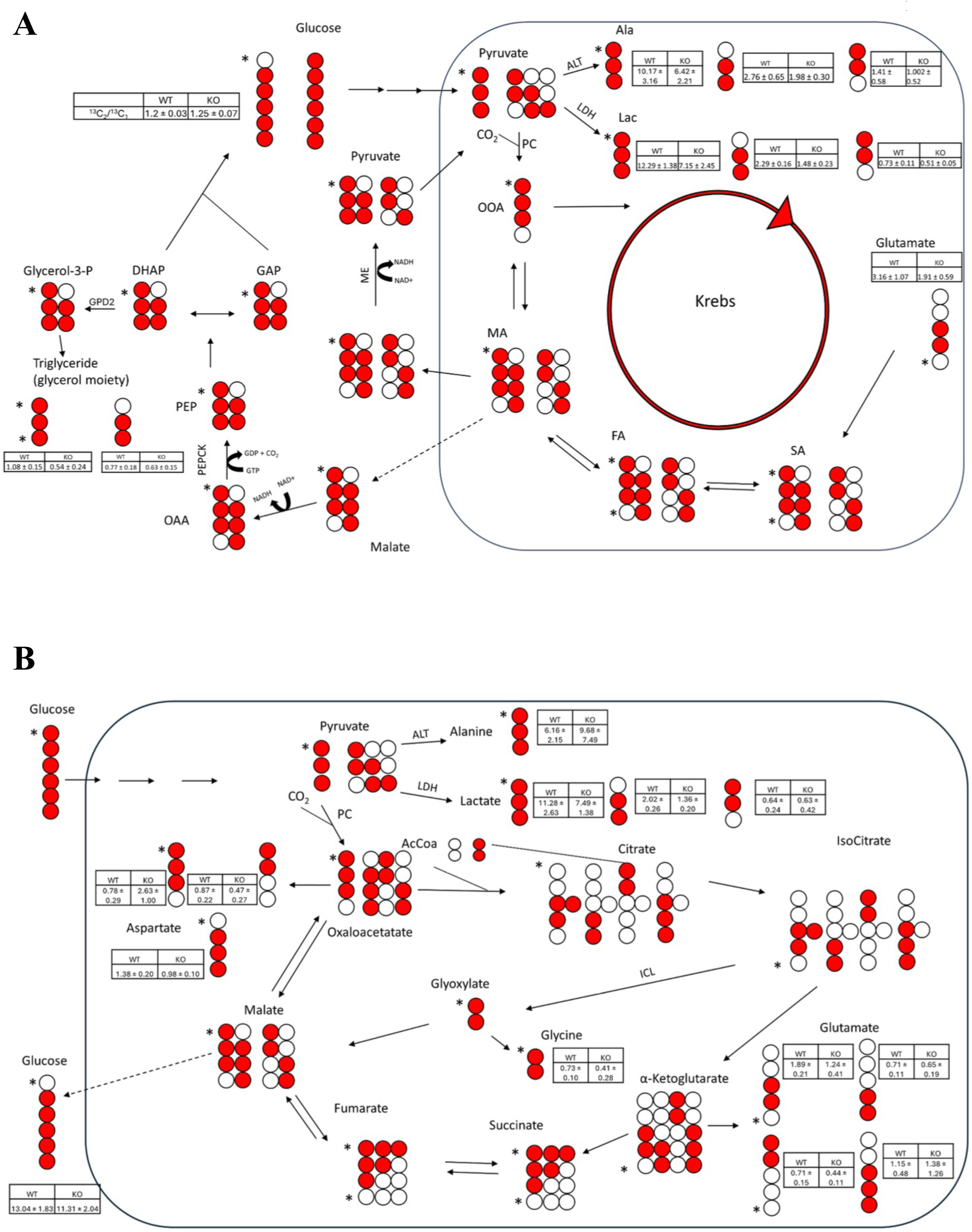
Schematic representation of proposed pathway for [1,2,3,4,5,6-^13^C_6_] glucose metabolism in liver (A) and kidney (B) from *Pde5a* ko and wild-type mice. Quantified species values are tabulated.

**Supplementary Figure 4.**
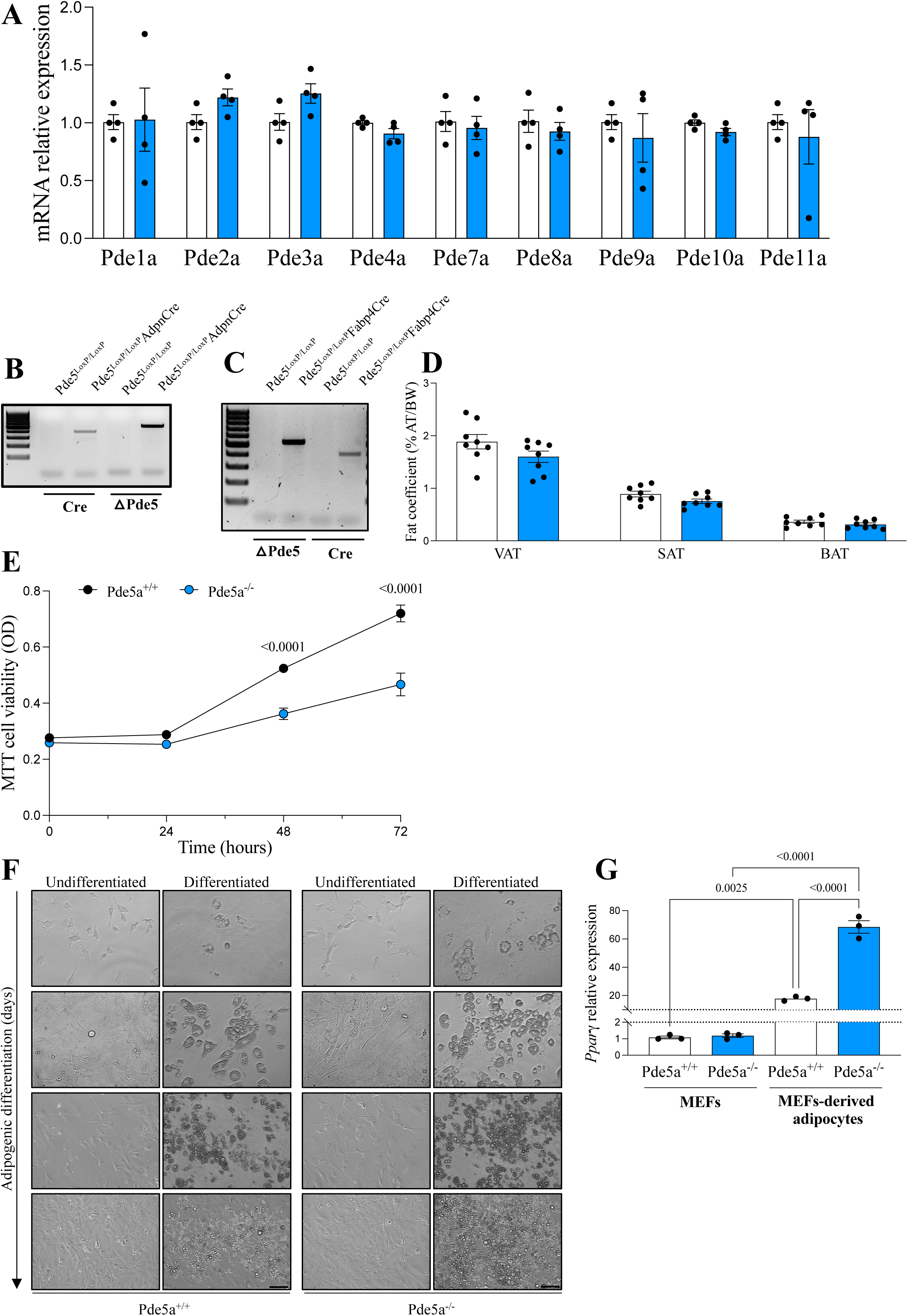
(A) qPCR gene expression analysis of *Pde1a, Pde2a, Pde3a, Pde4a, Pde7a, Pde8a, Pde9a, Pde10a* and *Pde11a* performed on visceral adipose tissue obtained from *Pde5a*^+/+^ (*n =4, white*) and *Pde5a*^-/-^ (*n =4, blue*) mice. *Hprt1* was used as housekeeping gene for normalization. Data are presented as dot plots with column bars ± SEM. Statistical analysis was performed using Student t-test. (B) Genotyping on genomic DNA isolated from tails of *Pde5a^LoxP/LoxP^ and Pde5a ^LoxP/LoxP^; Adpn^Cre^* showing a 432bp band corresponding to CRE and a 646bp band corresponding to *Pde5a* deleted allele *(ΔPde5a*). (C) Genotyping on genomic DNA isolated from tails of *Pde5a^LoxP/LoxP^ and Pde5a ^LoxP/LoxP^; Fabp4^Cre^* showing a 432bp band corresponding to CRE and a 646bp band corresponding to *Pde5a* deleted allele *(ΔPde5a*). (D) Adipose tissue (visceral adipose tissue VAT, subcutaneous adipose tissue SAT, brown adipose tissue BAT) weight as a percentage of total body weight of from *Pde5a^LoxP/LoxP^* (*white,* n=8) and *Pde5a ^LoxP/LoxP^; Fabp4^Cre^* (*blue,* n=8). Data are presented as dot plots with column bars ± SEM. Statistical analysis was performed using two-way ANOVA test. (E) MTT assay of MEF cells isolated from *Pde5a*^+/+^ (*n =3, white*) and *Pde5a*^-/-^ (*n=3, blue*) mice and cultured up to 96 hours. Data are presented as mean of optical density values ± SEM. Statistical analysis was performed using Student t-test. (F) Representative brightfield images of undifferentiated MEFs and MEF following differentiation into adipocytes at days 0, 5, 7 and 10. For differentiation medium composition see materials and methods section. Scale bar = 20 μm. (G) qPCR gene expression analysis of *Pparγ* performed on MEFs and MEFs-derived adipocytes obtained from *Pde5a*^-/-^ (*blue*) and wild-type (*white*) mice. *Hprt1* was used as housekeeping gene for normalization. Data are presented as dot plots with column bars ± SEM (n=3). Statistical analysis was performed using two-way ANOVA test.

**Supplementary Figure 5.**
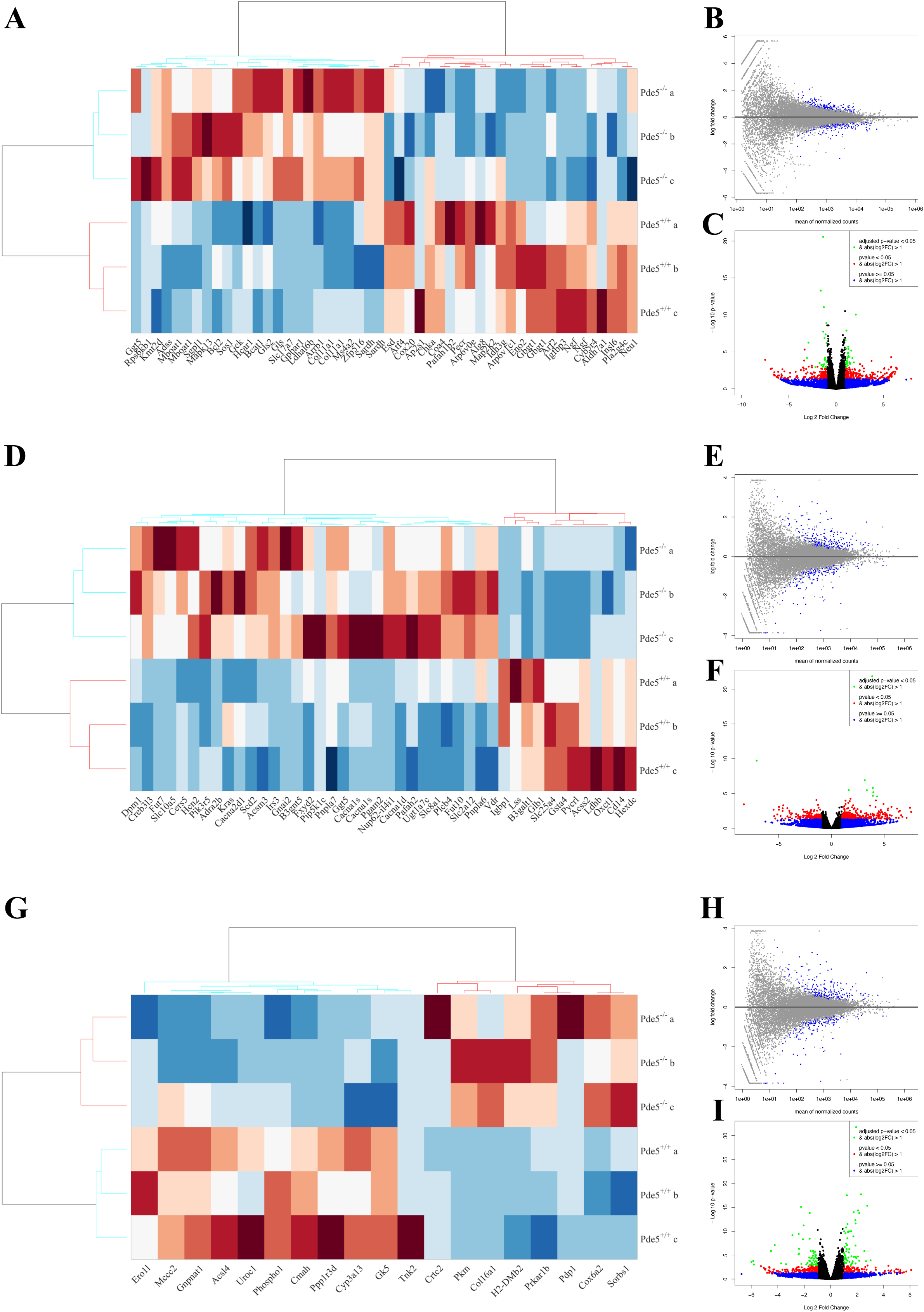
(A, D, G) A two-way hierarchical clustering dendrogram of DEseq genes in VAT (A), BAT (D) and liver (G) from *Pde5a* ko and wt mice (n=3). Data are expressed as mean FPKM, standardized, and visualized using the MATLAB “clustergram” script. Red: relatively high expression; blue: relatively low expression. (B, E, H) Mean average plot for VAT (B), BAT (E) and liver (H) showing genes (dots) with mean expression levels across all samples plotted on x-axis and log2 fold change observed in the contrast of interest on y-axis. Significant differentially expressed genes with p≤0.05 are colored in blue. (C, F, I) Volcano plot for VAT (C), BAT (F) and liver (I) comparing the amount of gene expression change (plotted as log2 fold change) to the significance of that change (plotted as the -log10 transformation of the multiple test adjusted *P* value), with each dot representing a single gene.

## REFERENCES

1 Zechner, R. et al. FAT SIGNALS--lipases and lipolysis in lipid metabolism and signaling. Cell Metab 15, 279–291 (2012). 10.1016/j.cmet.2011.12.018

2 Gesta, S., Tseng, Y. H. & Kahn, C. R. Developmental origin of fat: tracking obesity to its source. Cell 131, 242–256 (2007). 10.1016/j.cell.2007.10.004

3 Riazi, K. et al. The prevalence and incidence of NAFLD worldwide: a systematic review and meta-analysis. Lancet Gastroenterol Hepatol 7, 851–861 (2022). 10.1016/S2468-1253(22)00165-0

4 Campolo, F. et al. Identification of murine phosphodiesterase 5A isoforms and their functional characterization in HL-1 cardiac cell line. J Cell Physiol 233, 325–337 (2018). 10.1002/jcp.25880

5 Li, S. et al. Sildenafil induces browning of subcutaneous white adipose tissue in overweight adults. Metabolism 78, 106–117 (2018). 10.1016/j.metabol.2017.09.008

6 Mitschke, M. M. et al. Increased cGMP promotes healthy expansion and browning of white adipose tissue. FASEB J 27, 1621–1630 (2013). 10.1096/fj.12-221580

7 Zhang, X. et al. Sildenafil promotes adipogenesis through a PKG pathway. Biochem Biophys Res Commun 396, 1054–1059 (2010). 10.1016/j.bbrc.2010.05.064

8 Armani, A., Marzolla, V., Rosano, G. M., Fabbri, A. & Caprio, M. Phosphodiesterase type 5 (PDE5) in the adipocyte: a novel player in fat metabolism? Trends Endocrinol Metab 22, 404–411 (2011). 10.1016/j.tem.2011.05.004

9 Campolo, F., Pofi, R., Venneri, M. A. & Isidori, A. M. Priming metabolism with the type 5 phosphodiesterase: the role of cGMP-hydrolyzing enzymes. Curr Opin Pharmacol 60, 298–305 (2021). 10.1016/j.coph.2021.08.007

10 Pofi, R. et al. Sex-specific effects of daily tadalafil on diabetic heart kinetics in RECOGITO, a randomized, double-blind, placebo-controlled trial. Sci Transl Med 14, eabl8503 (2022). 10.1126/scitranslmed.abl8503

11 West, T. M. et al. Phosphodiesterase 5 Associates With beta2 Adrenergic Receptor to Modulate Cardiac Function in Type 2 Diabetic Hearts. J Am Heart Assoc 8, e012273 (2019). 10.1161/jaha.119.012273

12 Venneri, M. A. et al. PDE5 Inhibition Stimulates Tie2-Expressing Monocytes and Angiopoietin-1 Restoring Angiogenic Homeostasis in Diabetes. J Clin Endocrinol Metab 104, 2623–2636 (2019). 10.1210/jc.2018-02525

13 Fiore, D. et al. PDE5 Inhibition Ameliorates Visceral Adiposity Targeting the miR-22/SIRT1 Pathway: Evidence From the CECSID Trial. J Clin Endocrinol Metab 101, 1525–1534 (2016). 10.1210/jc.2015-4252

14 Giannetta, E. et al. Chronic Inhibition of cGMP phosphodiesterase 5A improves diabetic cardiomyopathy: a randomized, controlled clinical trial using magnetic resonance imaging with myocardial tagging. Circulation 125, 2323–2333 (2012). 10.1161/circulationaha.111.063412

15 Haas, B. et al. Protein kinase G controls brown fat cell differentiation and mitochondrial biogenesis. Sci Signal 2, ra78 (2009). 10.1126/scisignal.2000511

16 Ceddia, R. P. et al. Increased Energy Expenditure and Protection From Diet-Induced Obesity in Mice Lacking the cGMP-Specific Phosphodiesterase PDE9. Diabetes 70, 2823–2836 (2021). 10.2337/db21-0100

17 Moro, C., Klimcakova, E., Lafontan, M., Berlan, M. & Galitzky, J. Phosphodiesterase-5A and neutral endopeptidase activities in human adipocytes do not control atrial natriuretic peptide-mediated lipolysis. Br J Pharmacol 152, 1102–1110 (2007). 10.1038/sj.bjp.0707485

18 Cohen, P. & Kajimura, S. The cellular and functional complexity of thermogenic fat. Nat Rev Mol Cell Biol 22, 393–409 (2021). 10.1038/s41580-021-00350-0

19 Chouchani, E. T., Kazak, L. & Spiegelman, B. M. New Advances in Adaptive Thermogenesis: UCP1 and Beyond. Cell Metab 29, 27–37 (2019). 10.1016/j.cmet.2018.11.002

20 Haigh, J. L., New, L. E. & Filippi, B. M. Mitochondrial Dynamics in the Brain Are Associated With Feeding, Glucose Homeostasis, and Whole-Body Metabolism. Front Endocrinol (Lausanne) 11, 580879 (2020). 10.3389/fendo.2020.580879

21 Gu, X., Ma, Y., Liu, Y. & Wan, Q. Measurement of mitochondrial respiration in adherent cells by Seahorse XF96 Cell Mito Stress Test. STAR Protoc 2, 100245 (2021). 10.1016/j.xpro.2020.100245

22 Crane, J. D., Mottillo, E. P., Farncombe, T. H., Morrison, K. M. & Steinberg, G. R. A standardized infrared imaging technique that specifically detects UCP1-mediated thermogenesis in vivo. Mol Metab 3, 490–494 (2014). 10.1016/j.molmet.2014.04.007

23 Montgomery, M. K. et al. Mouse strain-dependent variation in obesity and glucose homeostasis in response to high-fat feeding. Diabetologia 56, 1129–1139 (2013). 10.1007/s00125-013-2846-8

24 Eccleston, H. B. et al. Chronic exposure to a high-fat diet induces hepatic steatosis, impairs nitric oxide bioavailability, and modifies the mitochondrial proteome in mice. Antioxid Redox Signal 15, 447–459 (2011). 10.1089/ars.2010.3395

25 Sun, Y. et al. High-fat diet promotes renal injury by inducing oxidative stress and mitochondrial dysfunction. Cell Death Dis 11, 914 (2020). 10.1038/s41419-020-03122-4

26 Kim, J., Oh, C. M. & Kim, H. The Interplay of Adipokines and Pancreatic Beta Cells in Metabolic Regulation and Diabetes. Biomedicines 11 (2023). 10.3390/biomedicines11092589

27 Tilg, H. & Moschen, A. R. Adipocytokines: mediators linking adipose tissue, inflammation and immunity. Nat Rev Immunol 6, 772–783 (2006). 10.1038/nri1937

28 Seo, D. H. et al. Effects of a Phosphodiesterase inhibitor on the Browning of Adipose Tissue in Mice. Biomedicines 10 (2022). 10.3390/biomedicines10081852

29 Reinhardt, R. R. et al. Distinctive anatomical patterns of gene expression for cGMP-inhibited cyclic nucleotide phosphodiesterases. J Clin Invest 95, 1528–1538 (1995). 10.1172/jci117825

30 Choi, Y. H. et al. Alterations in regulation of energy homeostasis in cyclic nucleotide phosphodiesterase 3B-null mice. J Clin Invest 116, 3240–3251 (2006). 10.1172/jci24867

31 Chung, Y. W. et al. White to beige conversion in PDE3B KO adipose tissue through activation of AMPK signaling and mitochondrial function. Sci Rep 7, 40445 (2017). 10.1038/srep40445

32 Zaccolo, M. & Movsesian, M. A. cAMP and cGMP signaling cross-talk: role of phosphodiesterases and implications for cardiac pathophysiology. Circ Res 100, 1569–1578 (2007). 10.1161/circresaha.106.144501

33 Honnor, R. C., Dhillon, G. S. & Londos, C. cAMP-dependent protein kinase and lipolysis in rat adipocytes. II. Definition of steady-state relationship with lipolytic and antilipolytic modulators. J Biol Chem 260, 15130–15138 (1985).

34 Djouder, N. et al. PKA phosphorylates and inactivates AMPKalpha to promote efficient lipolysis. EMBO J 29, 469–481 (2010). 10.1038/emboj.2009.339

35 Allen, M. D. & Zhang, J. Subcellular dynamics of protein kinase A activity visualized by FRET-based reporters. Biochem Biophys Res Commun 348, 716–721 (2006). 10.1016/j.bbrc.2006.07.136

36 Eguchi, J. et al. Transcriptional control of adipose lipid handling by IRF4. Cell Metab 13, 249–259 (2011). 10.1016/j.cmet.2011.02.005

37 Palmisano, B. et al. Gsalpha(R201C) and estrogen reveal different subsets of bone marrow adiponectin expressing osteogenic cells. Bone Res 10, 50 (2022). 10.1038/s41413-022-00220-1

38 Zhu, B., Strada, S. & Stevens, T. Cyclic GMP-specific phosphodiesterase 5 regulates growth and apoptosis in pulmonary endothelial cells. Am J Physiol Lung Cell Mol Physiol 289, L196–206 (2005). 10.1152/ajplung.00433.2004

39 Gebska, M. A. et al. Phosphodiesterase-5A (PDE5A) is localized to the endothelial caveolae and modulates NOS3 activity. Cardiovasc Res 90, 353–363 (2011). 10.1093/cvr/cvq410

40 Jeffery, E. et al. Characterization of Cre recombinase models for the study of adipose tissue. Adipocyte 3, 206–211 (2014). 10.4161/adip.29674

41 Dastagir, K. et al. Murine embryonic fibroblast cell lines differentiate into three mesenchymal lineages to different extents: new models to investigate differentiation processes. Cell Reprogram 16, 241–252 (2014). 10.1089/cell.2014.0005

42 Kim, M. S. et al. Association of genetic risk, lifestyle, and their interaction with obesity and obesity-related morbidities. Cell Metab 36, 1494–1503 e1493 (2024). 10.1016/j.cmet.2024.06.004

43 Lim, S. et al. Cold-induced activation of brown adipose tissue and adipose angiogenesis in mice. Nat Protoc 7, 606–615 (2012). 10.1038/nprot.2012.013

44 Wu, J. et al. Beige adipocytes are a distinct type of thermogenic fat cell in mouse and human. Cell 150, 366–376 (2012). 10.1016/j.cell.2012.05.016

45 Joseph, J. J. et al. Association of Adiposity With Incident Diabetes Among Black Adults in the Jackson Heart Study. J Am Heart Assoc 10, e020716 (2021). 10.1161/jaha.120.020716

46 Nedungadi, D. et al. The Association of Adiposity and RAAS with Incident Diabetes in African Americans: The Jackson Heart Study. J Clin Endocrinol Metab (2024). 10.1210/clinem/dgae396

47 Kowalski, G. M. & Bruce, C. R. The regulation of glucose metabolism: implications and considerations for the assessment of glucose homeostasis in rodents. Am J Physiol Endocrinol Metab 307, E859–871 (2014). 10.1152/ajpendo.00165.2014

48 Garcia-Villafranca, J., Guillen, A. & Castro, J. Involvement of nitric oxide/cyclic GMP signaling pathway in the regulation of fatty acid metabolism in rat hepatocytes. Biochem Pharmacol 65, 807–812 (2003). 10.1016/s0006-2952(02)01623-4

49 McGarry, J. D. & Foster, D. W. Regulation of hepatic fatty acid oxidation and ketone body production. Annu Rev Biochem 49, 395–420 (1980). 10.1146/annurev.bi.49.070180.002143

50 Li, Y. et al. AMPK phosphorylates and inhibits SREBP activity to attenuate hepatic steatosis and atherosclerosis in diet-induced insulin-resistant mice. Cell Metab 13, 376–388 (2011). 10.1016/j.cmet.2011.03.009

51 Guilherme, A. et al. Acetyl-CoA carboxylase 1 is a suppressor of the adipocyte thermogenic program. Cell Rep 42, 112488 (2023). 10.1016/j.celrep.2023.112488

52 Zhang, X., Yang, S., Chen, J. & Su, Z. Unraveling the Regulation of Hepatic Gluconeogenesis. Front Endocrinol (Lausanne) 9, 802 (2018). 10.3389/fendo.2018.00802

53 Watanabe, M. et al. Bile acids induce energy expenditure by promoting intracellular thyroid hormone activation. Nature 439, 484–489 (2006). 10.1038/nature04330

54 Kida, T., Tsubosaka, Y., Hori, M., Ozaki, H. & Murata, T. Bile acid receptor TGR5 agonism induces NO production and reduces monocyte adhesion in vascular endothelial cells. Arterioscler Thromb Vasc Biol 33, 1663–1669 (2013). 10.1161/atvbaha.113.301565

55 Kim, N. J. et al. A PDE1 inhibitor reduces adipogenesis in mice via regulation of lipolysis and adipogenic cell signaling. Exp Mol Med 51, 1–15 (2019). 10.1038/s12276-018-0198-7

56 Hankir, M. K. et al. A novel thermoregulatory role for PDE10A in mouse and human adipocytes. EMBO Mol Med 8, 796–812 (2016). 10.15252/emmm.201506085

57 Tomaszewski, M. R. et al. Magnetic resonance imaging detects white adipose tissue beiging in mice following PDE10A inhibitor treatment. J Lipid Res 64, 100408 (2023). 10.1016/j.jlr.2023.100408

58 Zhang, R., Maratos-Flier, E. & Flier, J. S. Reduced adiposity and high-fat diet-induced adipose inflammation in mice deficient for phosphodiesterase 4B. Endocrinology 150, 3076–3082 (2009). 10.1210/en.2009-0108

59 Guirguis, E. et al. A role for phosphodiesterase 3B in acquisition of brown fat characteristics by white adipose tissue in male mice. Endocrinology 154, 3152–3167 (2013). 10.1210/en.2012-2185

60 Fryk, E. et al. Feasibility of high-dose tadalafil and effects on insulin resistance in well-controlled patients with type 2 diabetes (MAKROTAD): a single-centre, double-blind, randomised, placebo-controlled, cross-over phase 2 trial. EClinicalMedicine 59, 101985 (2023). 10.1016/j.eclinm.2023.101985

61 Lopez-Varela, S., Sanchez-Muniz, F. J. & Cuesta, C. Decreased food efficiency ratio, growth retardation and changes in liver fatty acid composition in rats consuming thermally oxidized and polymerized sunflower oil used for frying. Food Chem Toxicol 33, 181–189 (1995). 10.1016/0278-6915(94)00133-9

62 Heine, M. et al. Lipolysis Triggers a Systemic Insulin Response Essential for Efficient Energy Replenishment of Activated Brown Adipose Tissue in Mice. Cell Metab 28, 644–655 e644 (2018). 10.1016/j.cmet.2018.06.020

63 Meyer, C. W., Ootsuka, Y. & Romanovsky, A. A. Body Temperature Measurements for Metabolic Phenotyping in Mice. Front Physiol 8, 520 (2017). 10.3389/fphys.2017.00520

64 Campolo, F. et al. cAMP-specific phosphodiesterase 8A and 8B isoforms are differentially expressed in human testis and Leydig cell tumor. Front Endocrinol (Lausanne) 13, 1010924 (2022). 10.3389/fendo.2022.1010924

65 Klein, S., Staring, M., Murphy, K., Viergever, M. A. & Pluim, J. P. elastix: a toolbox for intensity-based medical image registration. IEEE Trans Med Imaging 29, 196–205 (2010). 10.1109/tmi.2009.2035616

66 Monti, S. et al. RESUME(N): A flexible class of multi-parameter qMRI protocols. Phys Med 88, 23–36 (2021). 10.1016/j.ejmp.2021.04.005

67 Brancato, V. et al. Evaluation of a Whole-Liver Dixon-Based MRI Approach for Quantification of Liver Fat in Patients with Type 2 Diabetes Treated with Two Isocaloric Different Diets. Diagnostics (Basel) 12 (2022). 10.3390/diagnostics12020514

68 Vinue, A. & Gonzalez-Navarro, H. Glucose and Insulin Tolerance Tests in the Mouse. Methods Mol Biol 1339, 247–254 (2015). 10.1007/978-1-4939-2929-0_17

69 Barbagallo, F. et al. Genetically Encoded Biosensors Reveal PKA Hyperphosphorylation on the Myofilaments in Rabbit Heart Failure. Circ Res 119, 931–943 (2016). 10.1161/circresaha.116.308964

70 Errico, F. et al. The GTP-binding protein Rhes modulates dopamine signalling in striatal medium spiny neurons. Mol Cell Neurosci 37, 335–345 (2008). 10.1016/j.mcn.2007.10.007

71 Rosito, M. et al. Antibiotics treatment promotes vasculogenesis in the brain of glioma-bearing mice. Cell Death Dis 15, 210 (2024). 10.1038/s41419-024-06578-w

72 Miccheli, A. et al. Metabolic profiling by 13C-NMR spectroscopy: [1,2-13C2]glucose reveals a heterogeneous metabolism in human leukemia T cells. Biochimie 88, 437–448 (2006). 10.1016/j.biochi.2005.10.004

73 Gorietti, D. et al. 13C NMR based profiling unveils different alpha-ketoglutarate pools involved into glutamate and lysine synthesis in the milk yeast Kluyveromyces lactis. Biochim Biophys Acta 1850, 2222–2227 (2015). 10.1016/j.bbagen.2015.07.008

